# From Problems to Solutions: How Self-Questions Drive Strong Scientific Reasoning

**DOI:** 10.64898/2026.10.30.762163

**Authors:** Noelle Clark, Todd Montgomery, Gheed Nafea, Temilope Aina, Julie Dangremond Stanton

## Abstract

Metacognition, or awareness and control of thinking, is essential for problem solving because it helps solvers select strategies for learning and decide when to enact them (planning), check their understanding in the moment (monitoring), and assess effectiveness of solutions and learning strategies (evaluating). Most studies that examine metacognition do so retrospectively, leaving a critical gap in how metacognition is used in-the-moment for problem solving. In this study, we utilized a think-aloud interview approach to capture and characterize the metacognitive skills that senior undergraduate life science students at three institution types in the United States used spontaneously while solving challenging biology problems. We found that the students (n = 39) displayed a wide variety and quantity of monitoring, some evaluating, and nearly no planning skills while creating a solution backed with high-quality reasoning. Students frequently used self-questions to monitor and evaluate their thinking. We found that self-questioning is a critical skill for generating solutions, but it only moves problem solving forward if the student asks and correctly answers the self-question out loud. This study illustrates how self-questioning and other metacognitive regulation skills provide links to bridge the gap between a student’s prior knowledge and specific features of a challenging problem. Based on our student data, we conceptualized a model of metacognitive phases during problem solving with high-quality reasoning. Finally, we present a self-question scaffold to facilitate the proposed phases (planning, monitoring, reasoning, and evaluating) so educators can equip students with metacognitive and scientific reasoning tools for effective problem solving.

## INTRODUCTION

Problem solving is a crucial skill for students to develop in all scientific disciplines. Through the application of knowledge, problem solving facilitates students’ deep conceptual understanding (Frey et al., 2022). Metacognition, or the awareness and control of thinking for the purposes of learning (Stanton et al., 2021) is associated with improved problem-solving skills and performance (Coutinho et al., 2005; Goos et al., 2002; Ohtani & Hisasaka, 2018; Rickey & Stacy, 2000; Sandi-Urena et al., 2012). Problem solving and metacognition have been identified as important skills for students to develop in their science courses; however, the types of metacognition and how it facilitates successful scientific problem solving remains unclear. To bridge this knowledge gap, we investigated the metacognitive skills that fourth-year life science students use during problem solving to construct solutions backed with scientific reasoning. Based on our findings, we propose multiple implications for instructors to prompt and promote metacognitive and scientific reasoning skills during problem solving.

### Metacognition

Metacognition, or awareness and control of thinking, is imperative for learning and problem solving because it helps students gauge their level of understanding and take action for learning (Cross & Paris, 1988). In this study, we conceptualize metacognition based on Schraw and Moshman’s (1995) theoretical framework as opposed to Nelson and Narens’ (1990) framework. While Nelson and Narens (1990) focus specifically on the role of cognitive and metacognitive processes in memory acquisition, retention, and retrieval; Schraw and Moshman’s (1995) framework emphasizes the influence of metacognition on learning as a whole.

In order to broadly capture how individuals integrate metacognition into learning and knowledge construction, Schraw and Moshman’s theoretical framework divides metacognition into two facets: metacognitive knowledge and regulation (1995). Metacognitive knowledge refers to what we know about our process of learning, and the strategies that exist for learning. For instance, a student uses their metacognitive knowledge when they remember that if they read a question prompt only once, they often miss important information, so they always read it twice. Metacognitive regulation is how we regulate our thoughts while learning and the actions we take to learn (Sandi-Urena et al., 2012; Schraw & Moshman, 1995). For example, a student employs metacognitive regulation skills when they determine they understand a topic after applying their knowledge and solving a problem without referring to their notes. Then, they plan to do the same strategy for the next unit and adhere to their plan while learning.

In our study, we focus on metacognitive regulation, which can be broken down further into three skills: monitoring, planning, and evaluating. Students use these regulatory skills when they check understanding in the moment (monitoring), determine what to do in a future experience (planning), and assess effectiveness of a past plan and learning strategies (evaluating). For example, a student may monitor their understanding when they ask themself a question to confirm their knowledge while solving a challenging problem. They may evaluate if their strategy of using flashcards was effective based on a high score on their most recent exam; then, plan to use the same strategy to study for their next quiz. We choose to study metacognitive regulation because it includes skills that can be taught, learned, and developed over time (Rivas et al., 2022; Stanton et al., 2021).

### Metacognition and Problem Solving with High-Quality Reasoning

Metacognitive skill use is associated with improved and effective scientific problem-solving skills (Coutinho et al., 2005; Goos et al., 2002; Rickey & Stacy, 2000; Sandi-Urena et al., 2012). In order to solve scientific problems effectively, students must use high-quality reasoning (HQR) to support their claims. Informed by the Evidence-Based Reasoning framework proposed by Brown et al. 2010, we define HQR as the use of scientifically accurate and appropriate backing (data, evidence, and/or a rule) to link an argument premise to a claim. Take the following sentence for example, “The tree will sway because of the strong wind.” “The tree” is the premise, “will sway” is the claim, and, “because of the strong wind” is the backing. While this example illustrates the basic building blocks of constructing an argument, students must be able to expand on this framework in a scientific context, and create solutions to answer challenging problems.

While the ability to evaluate evidence and use reasoning to solve scientific problems begins to develop in adolescence (Vilma & Pillow, 2025), it is an integral skill that science students must continue developing in college (Bao et al., 2022). Efforts to reform undergraduate life science education have identified the ability to use HQR as a crucial competency for students to achieve during scientific training (AAAS, 2011). By nature, HQR is challenging for students; they must use their metacognitive skills to monitor their understanding and evaluate their learning strategies while they solve challenging science problems. When we push students to the bounds of their prior knowledge, metacognitive tools are more influential than intellectual ability in learning gains (Prins et al., 2006; Veenman & Spaans, 2005).

Even though metacognition and HQR are critical skills for life science students’ learning and success during college and their future careers, there has been little research examining the purpose of metacognition employed in domain-specific contexts, such as problem solving (Krieger et al., 2022). Our prior work suggests that engaging in group work promotes metacognition and that certain metacognitive skills, such as evaluating group members’ ideas, can lead to HQR (Halmo et al., 2022). However, outside of this previous research, the mechanism for how metacognitive regulation may contribute to HQR for life science problem solving remains largely underexplored.

It is possible that this gap in knowledge is yet to be filled because most studies that examine students’ metacognition do so retrospectively instead of in the moment. There are multiple reasons that researchers choose retrospective methods. First, it is easier and less time consuming to collect and analyze data versus in the moment methods. Additionally, retrospective methods, such as surveys, can be implemented in studies with large cohorts of participants (Veenman, 2011a). By expecting students to recall their thinking minutes or even days after they have completed a learning task, we may not capture all or even most of the metacognition that occurs at the moment of learning. Participants’ self-reported metacognitive skills and actual behavior may not be comparable, since recollection of earlier thought processes may be distorted by memory (Veenman, 2011a). Additionally, due to the complexity of metacognitive processes, collecting data after a task performance versus during may not allow us to measure the same skills (Saraç & Karakelle, 2012). Thus, in order to study the metacognition students use during problem solving, an in-the-moment data collection method is necessary.

### Present Study

In this study, we used a think-aloud approach to investigate in-the-moment metacognitive skills fourth-year life science students use to construct solutions with HQR during problem solving. In a think-aloud data collection method, students are instructed to share all of their thoughts out loud while working through the problem; which allows researchers to reliably access cognitive and metacognitive processes while students are solving the problems (Ericsson & Simon, 1980). We specifically examined fourth-year life science students as we reason they may show the greatest evidence of both metacognitive regulation and HQR, since they are at the culmination of their undergraduate degree. Ultimately, the mechanism for how metacognitive regulation may contribute to HQR during problem solving remains largely underexplored. Thus, we ask the following research questions:

1. What types of metacognitive regulation are associated with high-quality reasoning when fourth-year life science students solve problems?
2. How do fourth-year life science students use different types of metacognitive regulation to construct high-quality reasoning when solving challenging problems?

By characterizing the types of metacognitive statements and questions students use and how they use them to build solutions with HQR, we can provide implications for instructors on how to effectively cultivate students’ problem-solving skills. The knowledge obtained from answering these research questions will be used to foster science student’s metacognitive skills for greater learning, scientific reasoning, and performance in college and subsequent careers.

## METHODS

### Research Participants and Study Context

This study is one part of a 4-year longitudinal research study which aims to characterize life science students’ metacognitive skills and how they develop over time. Recruitment began in fall 2021 from in-person introductory biology and chemistry courses at three institutions in the Southeast United States: University of Georgia (UGA), University of North Georgia (UNG), and Georgia Gwinnett College (GGC). We strategically chose a range of institution types; UGA is a research institution, UNG is a master’s university, and GGC is a baccalaureate college. Students completed a consent form, the revised Metacognition Awareness Inventory (rMAI)(Harrison & Vallin, 2018), and they self-reported their age, gender, race, ethnicity, intended major, and first-generation status. We used a purposeful sampling method to select 52 life science students as participants that were representative of their institution’s demographics and displayed a range of rMAI scores. This paper focuses only on the data collected in the fourth year of the longitudinal study in which we had retained 39 of the original 52 participants. In the fourth year we had 19 participants from UGA, 15 from UNG, and 5 from GGC. We obtained written consent from all students to participant in the study which was reviewed and granted exempt status by the University of Georgia Institutional Review Board (# STUDY00006457) and the University of North Georgia Institutional Review Board (2021-003). All identifying information has been removed and all names in this paper are pseudonyms.

### Data Collection

We used a two-part qualitative interview approach that included both semi-structured and think-aloud interview portions. This study focuses on data from the think-aloud segment of the students’ 4^th^ year interview. Participants were contacted through their preferred form of communication (email or text) to schedule interviews. Interviews were conducted once a year between fall 2021 and spring 2025. Each semester, before their interview, the participants took a survey which included the rMAI (Harrison & Vallin, 2018), Self Efficacy for Learning and Performance subscale from the Motivated Strategies for Learning Questionnaire (Pintrich et al., 1993), and the Epistemic Belief Inventory (Schraw et al., 1995). All of the think-aloud interviews were conducted and recorded on Zoom and lasted 20-60 minutes. Including the semi-structured portion of the interview, the two-part interview process lasted up to from 1 hour and 30 mins to 2 hours and 15 minutes.

The participants were asked to “think-aloud” and solve two open-ended, published biochemistry problems (Bhatia et al., 2022; Halmo et al., 2018, 2020) and one genetics problem developed by our lab (see Supplementary Information). We conducted think-aloud interviews because this data collection method allows researchers to accurately access in the moment cognitive and metacognitive processes students are using during problem solving (Ericsson & Simon, 1980).

Interviewers made it explicit to the interviewees that we were not focused on the correctness of their answers and we were most interested in their thought processes. In the first, second, and third year, they were given a practice problem to accustom them to thinking aloud before solving the various life science problems. The first biochemistry problem, referred to as the “pathway flux problem”, involved predicting the flux of a disrupted metabolic pathway that had a component of negative feedback regulation (Bhatia et al., 2022). The other biochemistry problem, or the “protein X problem”, involved predicting non-covalent interactions and protein folding following an amino acid mutation (Halmo et al., 2018, 2020). The genetics problem, or the “pedigree problem”, involved predicting genotypes of autosomal recessive and X-linked dominant traits from a pedigree.

Every year of the study, students were given isomorphic versions of the pathway flux and the protein X problem, in a different order every time they interviewed. The pedigree problem was added in the fourth-year interviews to increase challenge and novelty, since students engage in metacognition when solving difficult problems (Al-Gaseem et al., 2020; Carr & Taasoobshirazi, 2008). If the participant fell silent for more than 5 seconds, interviewers prompted the students with “What are you thinking now?” and/or “Can you tell me more about that?” Since the interviews were conducted over Zoom, we asked them to type their written solutions into the chat and asked them to show any scratch paper to the camera that they may have used to jot notes or draw on. All interviews were transcribed by Rev.com or Temi.com and checked for accuracy by researchers in our lab.

### Scoring written and spoken solutions

We created exemplary solutions for each answer, which were used as a guide to create a scoring rubric for each question in the Y4 interviews. Two researchers (N.C. and T.A.) individually scored all of the participants’ written solutions and then discussed and arrived at consensus on any scores that were not in agreement. We used a one-way ANOVA test and found there were no significant differences in the cumulative scores and scores for each question between each institution (p > 0.01). The average performance score was 5.97(±3.96) out of 16 points total. Though not statistically significant, UGA had the highest mean performance (6.94±3.83), followed by UNG (5.26±4.30), then GGC (4.25±2.63).

### Primary analysis: evidence of metacognition and high-quality scientific reasoning

#### Coding for metacognitive utterances

We refined and updated an existing codebook (Halmo et al., 2024) to analyze the fourth-year interview transcripts (n = 39). Two researchers (N.C. and J.D.S.) open-coded a subset of fourth year transcripts and discussed any themes that emerged. The objective of open coding is for individual researchers to reflect on the contents and range of data and stay open to any ideas that may emerge from the data. We also wrote analytic memos throughout data analysis to document the coding and inquiry processes. Then, we used an elaborative coding method to build and refine existing our codebook, which was important because it was originally created for the first-year interview analysis (Halmo et al., 2024). Elaborative coding involves analyzing new data in a “top down” approach with an already established codebook (Auerbach & Silverstein, 2003; Saldaña, 2013). Through iterative discussion, existing code names were adjusted and new codes were added that aligned with emergent themes found in the fourth-year interviews. Then, the codebook was used to iteratively code all of the transcripts individually by one faculty member, one graduate student, and two undergraduate researchers (N.C., J.D.S., T.A., G.N.) with MaxQDA2020 software (VERBI Software; Berlin, Germany). The codebook was refined and updated throughout the coding process until the codebook was stabilized. All analyzed transcripts were coded to consensus by two or more researchers and any disagreements were resolved through discussion.

#### Coding for high-quality scientific reasoning

Since our lab has studied metacognition and scientific reasoning in group work problem solving contexts, we used a codebook previously developed in our lab as a starting point (Halmo et al., 2022). Three researchers. (N.C., T.M., and J.D.S.) developed a new codebook informed by Brown’s Evidence-Based Reasoning (EBR) Framework to help us identify HQR. Though the previous codebook was used in a group work problem solving context, it informed us how to identify individual components of reasoning units, or premise, claim, and backing in think aloud interview transcripts. Additionally, since the level of high-quality reasoning in the previous codebook was dependent on the nature of students’ interactions with each other, we removed this aspect and readjusted the levels of HQR to better fit the individual problem-solving data.

In this study we define HQR as including both complete and correct ideas. A complete and correct reasoning unit includes scientifically accurate premise, claim, and backing (Brown et al., 2010). Under the EBR framework there are different levels of backing quality; a rule is the highest quality, followed by evidence, and data. (Brown et al., 2010; Furtak et al., 2010). Data are discrete reports of past or present observations pertaining to the data figures provided in the problem in support of a claim (Brown et al., 2010). Evidence includes statements describing observed relationships in the data figures provided in the problem in support of a claim; they are less general than rules because they relate to a specific context (Brown et al., 2010). Finally, rules are statements describing a general relationship or principle in support of a claim (Brown et al., 2010). A correct reasoning unit only includes scientifically accurate ideas. If any incorrect ideas are introduced, they are corrected within the reasoning unit.

First, we identified individual components of reasoning units, or premise, claim, and backing in three participants’ fourth year interview transcripts. We then determined that the standard of reasoning unit completeness for the purpose of our scoring was the use of a claim and a type backing. We did not require the students to say the premise aloud, because the premise is given in the question. We also determined that students who shared solutions with only correct ideas were of higher quality than solutions that had mixed ideas (both correct and incorrect ideas that were never corrected). Next, we developed a scoring scheme in which the highest quality scientific reasoning includes solutions with only correct ideas, claim(s), and relevant backing(s). The types of backing also determined the score; students who used a rule as their backing in a complete and correct reasoning unit were given the highest score, followed by evidence, then data. For instance, in Table 1, Simone shows evidence of the highest quality reasoning out of the three participant examples because she uses correct claims backed by correct rules.

**Table 1.** Correct and complete high-quality reasoning examples from the protein X problem. The highest quality of backing is rule, followed by evidence, then data.

| **High-quality reasoning score** | **Description** | **Example** |
|---|---|---|
| Complete reasoning unit, backing with data, only correct ideas* | A reasoning unit where an individual student makes a claim which is supported by backing in the form of **data**, which are discrete reports of past or present observations pertaining to the data figures provided in the problem set | *“It's changing the amino acid and I see that it changed the whole chain and not just something very small. It all looks different to me so I would think that it would change [the noncovalent interactions]”* - Angelina |
| Complete reasoning unit, backing with evidence, only correct ideas* | A reasoning unit where an individual student makes a claim which is supported by backing in the form of **evidence**, which are statements describing observed relationships in the data figures provided in the problem set | *“So now we have oxygen and nitrogen, so there's a dipole moment and it's a very big change in structure. I think it would have an effect on the non-covalent interaction because before there is no dipole moment, but now there is a dipole moment; could be stronger in one direction than the other.”*- Erika |
| Complete reasoning unit, backing with rule, only correct ideas* (Highest quality of backing) | A reasoning unit where an individual student makes a claim which is supported by backing in the form of a **rule**, which are statements describing a general relationship or principle | *“Yes, there would be a change in the type of non-covalent interaction. Leucine is a nonpolar hydrophobic amino acid, whereas asparagine is a polar and hydrophilic amino acid. A new, a new non-covalent interaction would occur because asparagine also contains an amine and an oxygen that would be capable of H-bonding unlike leucine.”* - Simone |

\*If a student introduces an incorrect scientific idea, they correct it. All names are pseudonyms.

Two researchers (N.C. and T.M.) individually identified all reasoning components, determined accuracy of ideas included and then gave a score for the reasoning quality for each of the four solutions in all 39 fourth year interviews (i.e., 156 solutions total). We gave one reasoning quality score for the pathway flux problems, one score for the protein X problem, and two separate scores for the pedigree problem because there were two separate parts (A and B). Throughout this process, we regularly discussed and came to consensus regarding identifications of the reasoning components and scores of complete and correct reasoning units.

### Secondary analysis

#### Identifying what metacognitive utterances are associated with higher quality scientific reasoning

Two researchers (N.C. and T.M.) reviewed all transcript excerpts coded as having mixed ideas to identify student responses that contained more correct ideas than incorrect ideas. Mixed ideas included student solutions that contained nearly all correct ideas but had minor incorrect ideas. Additionally, mixed ideas also captured student solutions that contained nearly all incorrect ideas and limited correct ideas. This allowed us to include participant responses in our results who had nearly fully correct reasoning but provided a minor incorrect idea that didn’t impact the outcome of their final answer. We considered these responses HQR making note that the responses had mostly correct ideas instead of only correct ideas.

Then, we extracted the instances of HQR from MaxQDA and then identified the metacognitive codes that were prevalent in these problem-solving segments. Next, we systematically compiled a list of metacognitive codes associated with instances of HQR. Then, we went through the codes and documented the most prevalent themes. We also examined how any metacognitive skills identified were operationalized in each individual instance that a student used HQR and classified the utterance as productive or unproductive in the problem-solving process. A metacognitive utterance is **productive** if it actively advances problem solving and is **unproductive** if it does not lead to progress. As an example, a student may ask themselves a question (metacognitive utterance) in order to retrieve and say aloud relevant evidence to back up a correct claim. Thus, we recorded that the instance of self-questioning was productive since it led the student to a higher quality of reasoning and furthered their problem-solving process. We also identified if certain metacognitive utterances are less or not productive compared to others for facilitating the student’s HQR. Through this method we were able to distinguish which productive metacognitive utterances lead to HQR and how they are executed when life science students solve challenging problems. One author (N.C.) primarily completed secondary analysis while two authors (J.D.S. and T.M) reviewed the extracted lists and themes.

### Acknowledging assumptions of the think-aloud protocol

By collecting data with a think-aloud approach we operate under three main assumptions. The first assumption is that the students in this study have an internal monologue or speech that we can access when they verbally think aloud during problem solving. Inner speech can be defined as “the subjective experience of language in the absence of overt and audible articulation” (Alderson-Day & Fernyhough, 2015, p. 931). In psychology and cognitive sciences, inner speech has been investigated and theorized as being important for memory, self-regulation, and other complex cognitive processes such as logical reasoning (Alderson-Day & Fernyhough, 2015). Some studies estimate that inner speak occurs anywhere from 25 to 100% of waking life, while other researchers found that some individuals do not have any inner speech episodes at all (Alderson-Day & Fernyhough, 2015; Heavey & Hurlburt, 2008; Hurlburt et al., 2013; Klinger & Cox, 1987; Langland-Hassan, 2014). The wide range in estimates is likely due to the variability of inner speech across individuals (Alderson-Day & Fernyhough, 2015). Secondly, we assume some semblance of completeness, meaning that participants are verbally sharing as much of their inner speech as they are able. However, a limitation of this assumption is that not all instances of inner speech are conceived as words or can be verbalized into words (Alderson-Day & Fernyhough, 2015; Hurlburt et al., 2013). The third assumption is that inner speech is important for metacognition (Langland-Hassan, 2014). We operate under the assumption that the formulation of inner speech is what enables individuals to objectify and analyze their own thoughts in order to plan, monitor, and evaluate for purposes of learning and problem solving.

While acknowledging these assumptions, we emphasize that think-aloud protocols are an accurate way for researchers to reliably access participants’ cognitive and metacognitive processes (Ericsson & Simon, 1980).

## RESULTS

### What types of metacognitive regulation are associated with high-quality reasoning when fourth-year life science students solve problems?

When we asked fourth-year life science students to solve challenging biology problems out loud, we found mostly monitoring and some evaluating utterances (statements and questions) associated with HQR. We distinguished that productive monitoring utterances, or those that actively advance the problem-solving process, included monitoring understanding, monitoring familiarity, self-correction, acknowledging assumptions, acknowledging inferences and self-questions. The monitoring utterance types are described in detail in Table 2, and they are illuminated by examples from our student data.

**Table 2.**
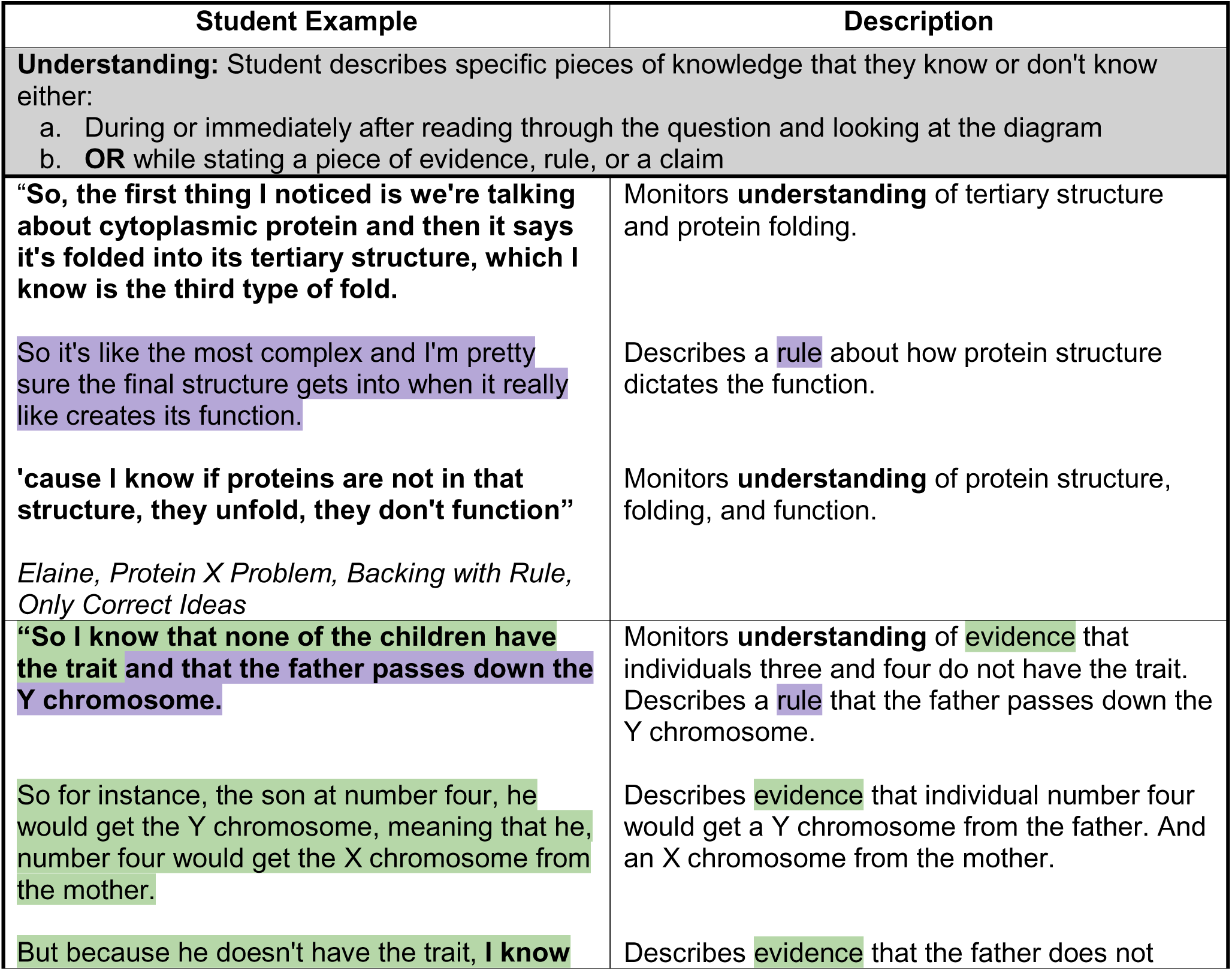
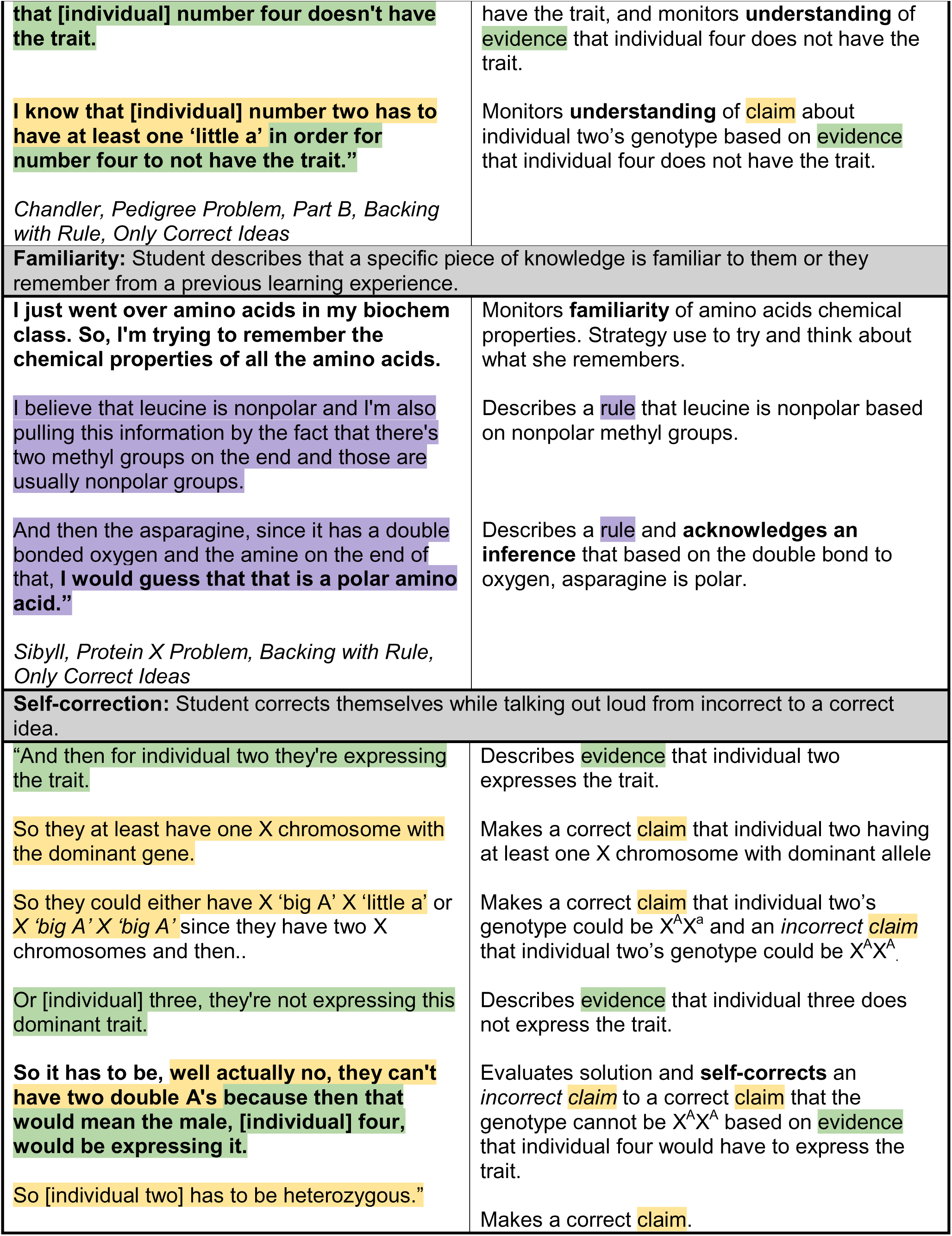
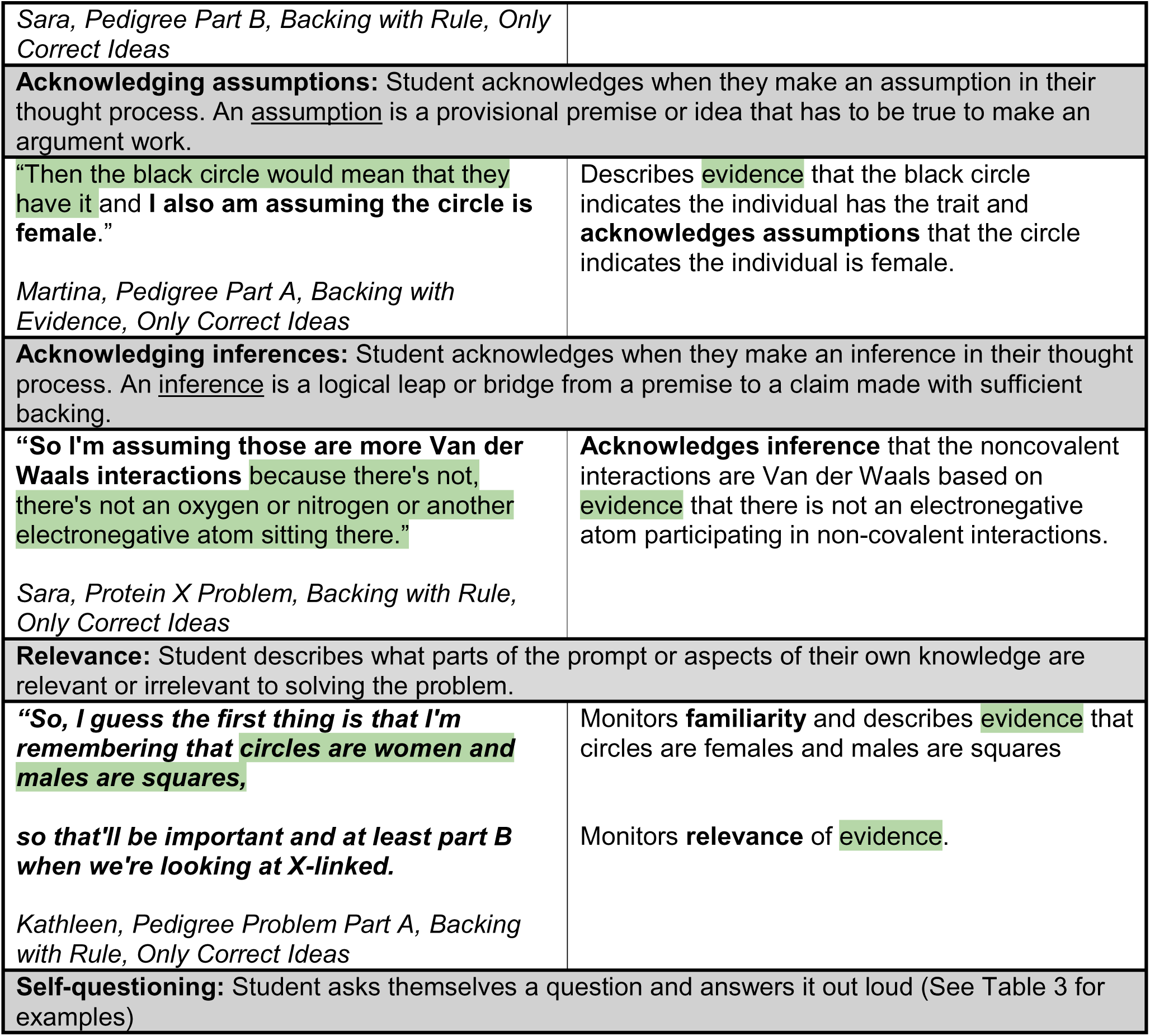
Monitoring utterances associated with high-quality reasoning. In the student example and description, the **monitoring utterances are bold** and evidence, rule, and claim are highlighted. All names are pseudonyms.

| Student Example | Description |
|---|---|
| **Understanding:** Student describes specific pieces of knowledge that they know or don't know either:<br>a. During or immediately after reading through the question and looking at the diagram<br>b. **OR** while stating a piece of evidence, rule, or a claim |  |
| “**So, the first thing I noticed is we're talking about cytoplasmic protein and then it says it's folded into its tertiary structure, which I know is the third type of fold.**<br>So it's like the most complex and I'm pretty sure the final structure gets into when it really like creates its function.<br>**'cause I know if proteins are not in that structure, they unfold, they don't function”**<br>*Elaine, Protein X Problem, Backing with Rule, Only Correct Ideas* | Monitors **understanding** of tertiary structure and protein folding.<br>Describes a rule about how protein structure dictates the function.<br>Monitors **understanding** of protein structure, folding, and function. |
| **“So I know that none of the children have the trait and that the father passes down the Y chromosome.**<br>So for instance, the son at number four, he would get the Y chromosome, meaning that he, number four would get the X chromosome from the mother.<br>But because he doesn't have the trait, **I know that [individual] number four doesn't have the trait.**<br>**I know that [individual] number two has to have at least one ‘little a’ in order for number four to not have the trait.”**<br>*Chandler, Pedigree Problem, Part B, Backing with Rule, Only Correct Ideas* | Monitors **understanding** of evidence that individuals three and four do not have the trait. Describes a rule that the father passes down the Y chromosome.<br>Describes evidence that individual number four would get a Y chromosome from the father. And an X chromosome from the mother.<br>Describes evidence that the father does not have the trait, and monitors **understanding** of evidence that individual four does not have the trait.<br>Monitors **understanding** of claim about individual two’s genotype based on evidence that individual four does not have the trait. |
| **Familiarity:** Student describes that a specific piece of knowledge is familiar to them or they remember from a previous learning experience. |  |
| **I just went over amino acids in my biochem class. So, I'm trying to remember the chemical properties of all the amino acids.**<br>I believe that leucine is nonpolar and I'm also pulling this information by the fact that there's two methyl groups on the end and those are usually nonpolar groups.<br>And then the asparagine, since it has a double bonded oxygen and the amine on the end of that, **I would guess that that is a polar amino acid.”**<br>*Sibyll, Protein X Problem, Backing with Rule, Only Correct Ideas* | Monitors **familiarity** of amino acids chemical properties. Strategy use to try and think about what she remembers.<br>Describes a rule that leucine is nonpolar based on nonpolar methyl groups.<br>Describes a rule and **acknowledges an inference** that based on the double bond to oxygen, asparagine is polar. |
| **Self-correction:** Student corrects themselves while talking out loud from incorrect to a correct idea. |  |
| “And then for individual two they're expressing the trait.<br>So they at least have one X chromosome with the dominant gene.<br>So they could either have X ‘big A’ X ‘little a’ or *X ‘big A’ X ‘big A’* since they have two X chromosomes and then..<br>Or [individual] three, they're not expressing this dominant trait.<br>**So it has to be, well actually no, they can't have two double A's because then that would mean the male, [individual] four, would be expressing it.**<br>So [individual two] has to be heterozygous.”<br>*Sara, Pedigree Part B, Backing with Rule, Only Correct Ideas* | Describes evidence that individual two expresses the trait.<br>Makes a correct claim that individual two having at least one X chromosome with dominant allele<br>Makes a correct claim that individual two’s genotype could be X<sup>A</sup>X<sup>a</sup> and an *incorrect* claim that individual two’s genotype could be X<sup>A</sup>X<sup>A</sup>.<br>Describes evidence that individual three does not express the trait.<br>Evaluates solution and **self-corrects** an *incorrect* claim to a correct claim that the genotype cannot be X<sup>A</sup>X<sup>A</sup> based on evidence that individual four would have to express the trait.<br>Makes a correct claim. |
| **Acknowledging assumptions:** Student acknowledges when they make an assumption in their thought process. An assumption is a provisional premise or idea that has to be true to make an argument work. |  |
| “Then the black circle would mean that they have it and **I also am assuming the circle is female**.”<br>*Martina, Pedigree Part A, Backing with Evidence, Only Correct Ideas* | Describes evidence that the black circle indicates the individual has the trait and **acknowledges assumptions** that the circle indicates the individual is female. |
| **Acknowledging inferences:** Student acknowledges when they make an inference in their thought process. An inference is a logical leap or bridge from a premise to a claim made with sufficient backing. |  |
| **“So I'm assuming those are more Van der Waals interactions** because there's not, there's not an oxygen or nitrogen or another electronegative atom sitting there.”<br>*Sara, Protein X Problem, Backing with Rule, Only Correct Ideas* | **Acknowledges inference** that the noncovalent interactions are Van der Waals based on evidence that there is not an electronegative atom participating in non-covalent interactions. |
| **Relevance:** Student describes what parts of the prompt or aspects of their own knowledge are relevant or irrelevant to solving the problem. |  |
| ***“So, I guess the first thing is that I'm remembering that circles are women and males are squares,***<br>***so that'll be important and at least part B when we're looking at X-linked.***<br>*Kathleen, Pedigree Problem Part A, Backing with Rule, Only Correct Ideas* | Monitors **familiarity** and describes evidence that circles are females and males are squares<br>Monitors **relevance** of evidence. |
| **Self-questioning:** Student asks themselves a question and answers it out loud (See Table 3 for examples) |  |

We found a variety of monitoring and some evaluating self-question types associated with HQR (Table 3 and 4). Self-questions, or asking yourself questions, were only productive and associated with HQR when the student answered their own question correctly out loud. In Table 3 we describe six categories of monitoring self-questions: self-questions to recall knowledge, identify problem features, confirm understanding, check accuracy, monitor conceptual understanding, and restating the problem in their own words (Table 3). Evaluative self-questions associated with HQR were far less common compared to monitoring self-questions associated with HQR. Students use evaluative self-questions to assess their work after a phase of problem-solving is complete. For example, students use evaluative self-questions after providing a claim to answer part of the problem or after they gave their full solution. In our data we found that students used evaluative self-questions including self-questions to assess accuracy, evaluate what the question is asking, and evaluate understanding (Table 4).

**Table 3.** Monitoring self-questioning types associated with high-quality reasoning. In the student example and description, the **monitoring utterances are bold** and evidence, rule, and claim are highlighted. Interestingly, self-questions were only associated with high-quality reasoning when students answered their own questions. All names are pseudonyms.

| Student Example | Description |
|---|---|
| **Recall knowledge:** Students ask themselves a question to remember a piece of relevant knowledge to solve the problem. Often, these questions start with the word “what.” The knowledge that they are seeking might be a word, a concept, a diagram component, or what they said or thought about previously. |  |
| **“Oh I, yeah, okay. I would need to draw the um; I forget what they're called. It's been so long.**<br>**Um, like draw out the interactions with the, what are they called?**<br>Well, I guess this is something I'd say I wish I could look up. Um, but the, like you draw the dots and then you count how they interact or whatever. So let me just think of what I remember though.<br>Um, oh, the valence electrons, that's what I'm talking about.”<br>*Elaine, Protein X Problem, Backing with Rule, Only Correct Ideas* | Monitors **familiarity.**<br>**Self-question to recall knowledge.**<br>Strategy use to try and think about what she remembers.<br>Answers self-question with recalled knowledge - valence electrons. |
| **“So [individual] one has to, since it's male and only has one X chromosome, it has to be…I think it's like… how is the subscription stuff to write this out?**<br>**I know it's X... On the X chromosome, it has a “little a” and on Y it doesn't really have (anything).”**<br>*Sara, Pedigree Problem Part A, Backing with Rule, Only Correct Ideas* | **Self-question to recall knowledge** of how to write a genotype<br>Monitors **understanding** of claim that individual one has the genotype X<sup>a</sup>Y. |
| **Identify problem features:** Students ask themselves a question to help them identify where something is in the diagram or to orient themselves to what they are looking at. |  |
| “**Okay, where is protein X?**<br>Okay. It's just the whole thing is folded into its tertiary structure surrounded by water molecules.”<br>*Jeffrey, Protein X Problem, Backing with Rule, Only Correct Ideas* | **Self-question to identify problem features.**<br>Answers self-question and describes where protein X is. |
| **Confirm understanding:** Students add a question inflection to a statement of recalled knowledge or a claim indicating that they are monitoring their understanding of what they just said. Nearly all of these examples, except one is from Simone’s interview. |  |
| **“Leucine is a nonpolar... Is nonpolar...?**<br>Polar can't cross membranes, which means it's hydrophilic. Alright.<br>**Polar is hydrophilic 'cause water?**<br>Yeah.<br>**So non-polar would be... leucine is nonpolar and hydrophobic?**<br>MmMm [affirmative]<br>Whereas asparagine is polar and hydrophilic.”<br>*Simone, Protein X Problem, Backing with Rule, Only Correct Ideas* | **Self-question to confirm understanding** that leucine is nonpolar.<br>Describes a rule about how polar molecules (generally) can’t cross cell membranes.<br>**Self-question to confirm understanding** that polar molecules are hydrophilic.<br>Answers self-question.<br>**Self-question to confirm understanding** of a rule that leucine is nonpolar and hydrophobic.<br>Answers self-question.<br>Describes a rule that asparagine is polar and hydrophilic. |
| **Check accuracy:** Students check the accuracy of a statement they just made, which happens in the moment. |  |
| “**Okay, so for [individual] one, so that's a male, right?**<br>I'm trying to remember. I think circles are female. Yeah. So [individual] one is a male.<br>So, [individual one] would have XY chromosomes.<br>And actually, let's skip [individual] one for now because that's more thinking than the other ones.”<br>*Jeffery, Pedigree Problem Part B, Backing with Rule, Only Correct Ideas* | **Self-question to check accuracy** that individual one is a male.<br>Describes evidence that circles are female and individual one is a male.<br>Describes rule that males have XY chromosomes.<br>Strategy that he is going to skip and come back to solve individual one’s genotype. |
| **Conceptual understanding:** Students ask themselves a conceptual question to monitor their understanding. |  |
| “Yes, there would be a change in the type of non-covalent interaction pointed to by the arrow.<br>**Why?**<br>Leucine is a nonpolar and hydrophobic amino acid whereas asparagine is a polar and hydrophilic amino acid.<br>[Asparagine] also contains an amine and an oxygen that would be capable of hydrogen bonding unlike leucine.”<br>*Simone, Protein X Problem, Backing with Rule, Only Correct Ideas* | Makes a claim that there would be a change in the non-covalent interaction.<br>**Conceptual understanding self-question.**<br>States rule that leucine is nonpolar and hydrophobic and asparagine is polar and hydrophilic.<br>States a rule that asparagine is capable of hydrogen bonding. |
| **Restating question in their own words** |  |
| **“Um, do we predict that [the mutation] would affect the non-covalent interaction?**<br>Yes, it would affect the non-covalent interaction, pointed to by the arrow.”<br>*Eva, Protein X Problem, Backing with Rule, Mostly Correct Ideas* | **Restates question in own words.**<br>States claim that the mutation would affect the noncovalent interaction. |

**Table 4.**
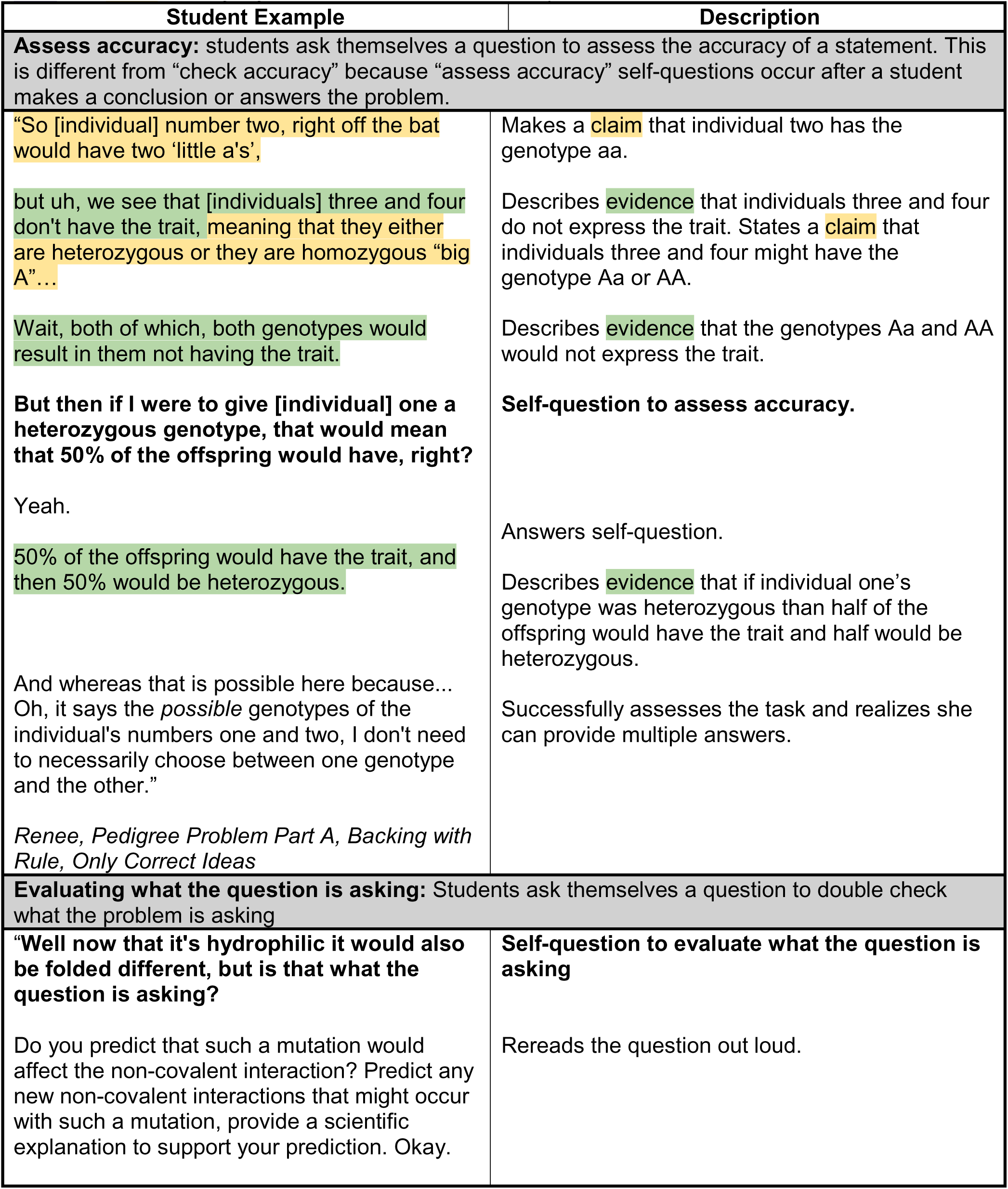
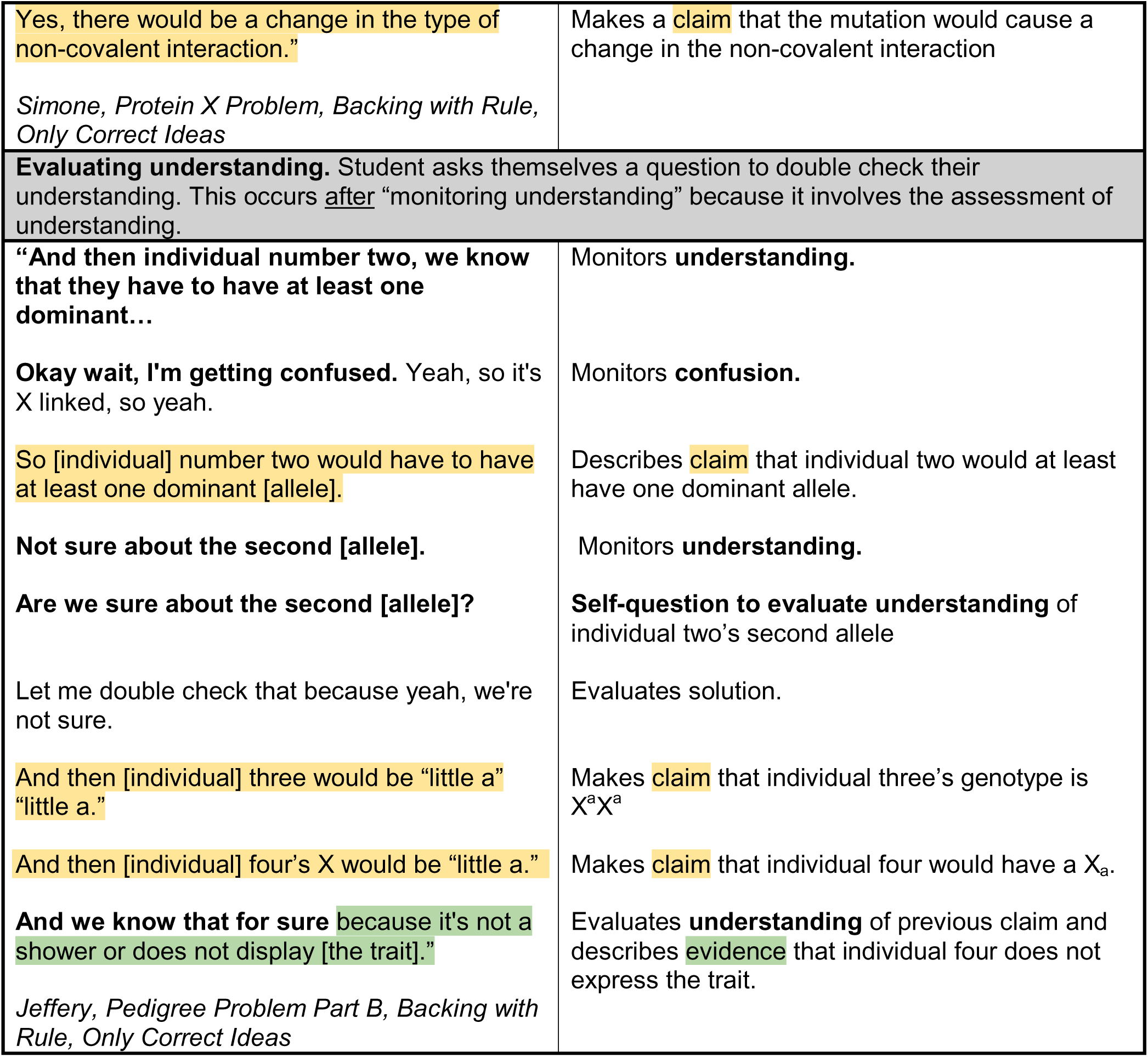
Evaluative self-questioning types associated with high-quality reasoning. In the student example and description, the **monitoring and evaluating utterances are bold** and evidence, rule, and claim are highlighted. All names are pseudonyms.

| Student Example | Description |
|---|---|
| **Assess accuracy:** students ask themselves a question to assess the accuracy of a statement. This is different from “check accuracy” because “assess accuracy” self-questions occur after a student makes a conclusion or answers the problem. |  |
| “So [individual] number two, right off the bat would have two ‘little a's’,<br>but uh, we see that [individuals] three and four don't have the trait, meaning that they either are heterozygous or they are homozygous “big A”…<br>Wait, both of which, both genotypes would result in them not having the trait.<br>**But then if I were to give [individual] one a heterozygous genotype, that would mean that 50% of the offspring would have, right?**<br>Yeah.<br>50% of the offspring would have the trait, and then 50% would be heterozygous.<br>And whereas that is possible here because... Oh, it says the *possible* genotypes of the individual's numbers one and two, I don't need to necessarily choose between one genotype and the other.”<br>*Renee, Pedigree Problem Part A, Backing with Rule, Only Correct Ideas* | Makes a claim that individual two has the genotype aa.<br>Describes evidence that individuals three and four do not express the trait. States a claim that individuals three and four might have the genotype Aa or AA.<br>Describes evidence that the genotypes Aa and AA would not express the trait.<br>**Self-question to assess accuracy.**<br>Answers self-question.<br>Describes evidence that if individual one’s genotype was heterozygous than half of the offspring would have the trait and half would be heterozygous.<br>Successfully assesses the task and realizes she can provide multiple answers. |
| **Evaluating what the question is asking:** Students ask themselves a question to double check what the problem is asking |  |
| “**Well now that it's hydrophilic it would also be folded different, but is that what the question is asking?**<br>Do you predict that such a mutation would affect the non-covalent interaction? Predict any new non-covalent interactions that might occur with such a mutation, provide a scientific explanation to support your prediction. Okay. | **Self-question to evaluate what the question is asking**<br>Rereads the question out loud. |
| Yes, there would be a change in the type of non-covalent interaction.”<br>*Simone, Protein X Problem, Backing with Rule, Only Correct Ideas* | Makes a claim that the mutation would cause a change in the non-covalent interaction |
| **Evaluating understanding.** Student asks themselves a question to double check their understanding. This occurs after “monitoring understanding” because it involves the assessment of understanding. |  |
| **“And then individual number two, we know that they have to have at least one dominant…**<br>**Okay wait, I'm getting confused.** Yeah, so it's X linked, so yeah.<br>So [individual] number two would have to have at least one dominant [allele].<br>**Not sure about the second [allele].**<br>**Are we sure about the second [allele]?**<br>Let me double check that because yeah, we're not sure.<br>And then [individual] three would be “little a” “little a.”<br>And then [individual] four’s X would be “little a.”<br>**And we know that for sure** because it's not a shower or does not display [the trait].”<br>*Jeffery, Pedigree Problem Part B, Backing with Rule, Only Correct Ideas* | Monitors **understanding.**<br>Monitors **confusion.**<br>Describes claim that individual two would at least have one dominant allele.<br>Monitors **understanding.**<br>**Self-question to evaluate understanding** of individual two’s second allele<br>Evaluates solution.<br>Makes claim that individual three’s genotype is X<sup>a</sup>X<sup>a</sup><br>Makes claim that individual four would have a X<sub>a</sub>.<br>Evaluates **understanding** of previous claim and describes evidence that individual four does not express the trait. |

### How do fourth-year life science students use different types of metacognitive regulation to construct high-quality reasoning when solving challenging problems?

Metacognitive regulation while problem solving (largely monitoring utterances, some evaluative self-questioning) provides links to bridge the gap between a student’s prior knowledge and specific features of a challenging problem. Below, we will describe how students use different types of monitoring and evaluating utterances that facilitate HQR. By identifying the purpose of each utterance type we were able to reveal that monitoring (checking understanding in the moment) aids students in the retrieval of knowledge, verification of knowledge accuracy and relevance, building of reasoning bridges, orienting themselves to what they are looking at, and reassessing what the question is asking. Evaluating (assessing effectiveness of solutions and learning strategies) aids students in double checking accuracy, understanding, and determining what the question is asking. Fourth-year life science students use a variety of metacognitive regulation skills, mostly monitoring and some evaluating utterances, to construct high-quality reasoning when solving challenging biology problems.

### Purpose of Monitoring Utterances

#### Retrieval of knowledge

Students in our study often needed to retrieve prior knowledge that they have acquired throughout their educational experiences to solve the challenging biology problems with HQR. We identified that students monitored understanding, familiarity, and asked self-questions to help retrieve their prior knowledge (Table 2).

##### Monitoring understanding to retrieve knowledge (Table 2)

Students monitored understanding when they describe pieces of knowledge that they do or do not know. This utterance was found across the data, regardless of reasoning quality. However, this utterance was associated with HQR when students monitored understanding immediately after reading the problem statement or while stating a piece of evidence, rule, or a claim.

For example, immediately after reading the protein X problem out loud, Elaine monitored understanding, *“So, the first thing I noticed is we’re talking about cytoplasmic protein and then it says it’s folded into its tertiary structure, which I know is the third type of fold.”* Then, she retrieves a rule, ‘*So it’s like the most complex and I’m pretty sure the final structure gets into when it really like creates its function.”* Finally, she monitors understanding again, “*cause I know if proteins are not in that structure, they unfold, they don’t function.”* Elaine proceeded to solve the problem correctly and used rules to back up her solution.

As another example, while solving the pedigree problem part B, Chandler monitored understanding while stating a piece of evidence, rule, or a claim. First, he monitored his understanding of evidence, followed by a rule, “*So I know that none of the children have the trait and that the father passes down the Y chromosome.”* Then, he retrieves evidence, *“So for instance, the son at number four, he would get the Y chromosome, meaning that he, number four would get the X chromosome from the mother.”* Next, he describes evidence and monitors understanding of evidence, “*But because he doesn’t have the trait, I know that [individual] number four doesn’t have the trait.”* Finally, Chandler monitors understanding of a claim, *“I know that [individual] number two has to have at least one ‘little a’”* which is backed by evidence, “*in order for number four to not have the trait.”* Chandler proceeded to solve the problem correctly and used a rule to back up his solution.

##### Monitoring familiarity to retrieve knowledge (Table 2)

Students monitor familiarity when they describe that a specific piece of knowledge is familiar or that they remember it from a previous learning experience. Monitoring familiarity was only productive and associated with HQR if the student described that a specific piece of knowledge is familiar to them or that they remember it from a previous learning experience (Table 2). Monitoring familiarity is not associated with HQR if a student described that a specific piece of knowledge is not familiar to them or that they forgot it from a previous learning experience. Interestingly, we found that monitoring familiarity was most productive when students used these utterances immediately after reading the question.

For example, after reading the question prompt, Sibyll monitored her familiarity and described a strategy of thinking about what she remembers, *“I just went over amino acids in my biochem class. So, I’m trying to remember the chemical properties of all the amino acids.”* Then, Sibyll retrieved a scientific rule from her memory, *“I believe that leucine is nonpolar and I’m also pulling this information by the fact that there’s two methyl groups on the end and those are usually nonpolar groups.”* Then she retrieved another rule and acknowledges an inference, *“And then the asparagine, since it has a double bonded oxygen and the amine on the end of that, I would guess that that is a polar amino acid.”* Then Sibyll proceeded to solve the problem correctly and used rules to back up her solution.

##### Monitoring with self-questions to retrieve knowledge (Table 3)

Students use self-questioning when they ask themselves a question and answer it out loud. There are two types of self-questions that serve the purpose of knowledge retrieval which are (1) self-questioning to recall knowledge and (2) self-questioning to monitor conceptual understanding.

Students used a self-question to recall knowledge when they asked themselves a question to remember a piece of relevant information to solve the problem. Often, these questions start with the word “what.” The knowledge that they are seeking might be a word, a concept, a component of a diagram, or what they said or thought about previously. For example, after monitoring that she forgot a piece of knowledge while solving the protein X problem, Elaine asked, *“Um, like draw out the interactions with the, what are they called?”* Then, she used a strategy to think about what she remembered and retrieved the knowledge to answer her own question, *“Um, oh, the valence electrons, that’s what I’m talking about.”*

As another example, while attempting to write a genotype while solving the pedigree part B problem, Sara asked herself, *“So [individual] one has to, since it’s male and only has one X chromosome, it has to be…I think it’s like… how is the subscription stuff to write this out?”* Then, Sara answered her question by monitoring her understanding of a claim, *“I know it’s X… On the X chromosome, it has a “little a” and on Y it doesn’t really have (anything).”* In the word choice of her self-question, Sara makes a small error, since X-linked dominant traits are written as superscripts and not subscripts. However, this mistake didn’t impact her conceptual understanding of the problem because she proceeded to solve the problem correctly using rules to back up her solution.

Some students also use self-questioning to monitor conceptual understanding while solving the problem. For example, Simone, stated a claim to answer the protein X problem, *“Yes, there would be a change in the type of non-covalent interaction pointed to by the arrow.”* Then, she asked herself a question to monitor her conceptual understanding, *“Why?”* She answered her self-question with two rules to back her claim, *“Leucine is a nonpolar and hydrophobic amino acid whereas asparagine is a polar and hydrophilic amino acid. [Asparagine] also contains an amine and an oxygen that would be capable of hydrogen bonding unlike leucine.”* After this example, Simone evaluated what the question is asking before submitting her complete and correct solution, with claims backed by rules.

#### Verification of knowledge accuracy and relevance

To solve questions effectively, students verify their knowledge accuracy and relevance during the problem-solving process. To verify knowledge accuracy and relevance, students use self-corrections, monitor relevance, and ask self-questions that can lead them to successfully solve problems with HQR (Tables 2).

##### Monitoring with self-correction to verify knowledge accuracy (Table 2)

Self-correction from an incorrect idea to a correct idea is a monitoring skill used by students to verify the accuracy of their thoughts, which can lead to HQR. While solving the pedigree problem part B, Sara made a correct claim, *“So [individual two] could either have X ‘big A’ X ‘little a’”*, followed by an incorrect claim, “*Or X ‘big A’ X ‘big A.’”* Then she describes a piece of evidence, *“Or [individual] three, they’re not expressing this dominant trait.”* Finally, she evaluates her incorrect solution and self-corrects the incorrect claim to a correct claim, *“So it has to be, well actually no, they can’t have two double A’s.”* The claim is then backed by evidence*, “Because then that would mean the male, [individual] four, would be expressing it.”* Then she described a correct claim, *“So [individual two] has to be heterozygous.”* Through monitoring by self-correction of an incorrect to a correct claim, Sara was able to solve the problem correctly, backing her claims with a rule.

##### Monitoring with self-questions to verify knowledge accuracy (Table 3)

Students can verify knowledge correctness with self-questions to confirm understanding and check accuracy, which are associated with HQR.

To confirm understanding and accuracy, students may add a question inflection to a statement of recalled knowledge or a claim indicating that they are monitoring their understanding of what they just said. Nearly all of these examples, except one is from Simone’s interview, highlighting how monitoring skills can be unique or even stylized across individuals. For example, Simone confirmed her understanding by describing *“Leucine is a nonpolar… Is nonpolar…?”* Then stated a rule, *“Polar can’t cross membranes, which means it’s hydrophilic. Alright.”* After, she asked herself another question to confirm understanding, *“Polar is hydrophilic ‘cause water?”* and answers *“Yeah.”* Finally, she asked herself another question to confirm understanding and said, *“So non-polar would be… leucine is nonpolar and hydrophobic?”* and answered “Mm-hm” affirmatively. Then, she described a rule, *“Whereas asparagine is polar and hydrophilic.”* As a result of monitoring to confirm understanding, Simone solved the protein X problem, backing her claim with rules.

Students also check the accuracy of a statement they just made by monitoring with a self-question to confirm accuracy. After reading the pedigree problem part B, Jeffery asked himself, *“Okay, so for [individual] one, so that’s a male, right?”* Then he confirmed his understanding, *“I’m trying to remember. I think circles are female. Yeah. So, [individual] one is a male.”* Then, he described a rule, *“So, [individual one] would have XY chromosomes.”* Jeffery then stated an interesting strategy, *“And actually, let’s skip [individual] one for now because that’s more thinking than the other ones.”* Jeffery then returned back to individual two’s genotype and provided a correct claim. Ultimately, Jeffery came to a solution with correct ideas about nearly all the individuals’ genotypes, but monitored that he did not know what the second allele of individual two’s genotype is.

##### Monitoring relevance to verify knowledge relevance (Table 2)

Students who successfully solve the problems with HQR occasionally monitor relevance, which is when a student describes what parts of the prompt or aspects of their own knowledge are relevant or irrelevant to solving the problem. Though not seen as frequently as other monitoring utterances, monitoring relevance is a crucial skill for solving problems with HQR because it can redirect students from going down an irrelevant train of thought. Monitoring relevance can also direct student thinking productively towards a successful solution. For example, immediately after reading the pedigree problem part A, Kathleen monitored familiarity of a piece of evidence, *“So, I guess the first thing is that I’m remembering that circles are females and males are squares.”* Then, she monitored the relevance of such evidence, “*So that’ll be important and at least part B when we’re looking at X-linked.”* Kathleen then goes on to successfully solve part A and B of the pedigree problem with HQR, backing claims with rules.

#### Building reasoning bridges

Students build reasoning bridges while solving problems when they apply their knowledge to make inferences and acknowledge underlying assumptions, which is associated with high-quality reasoning.

##### Acknowledging assumptions and inferences to build reasoning bridges (Table 2)

Students acknowledge assumptions when they name an assumption in their thought process. In the context of this study, an assumption is a premise or idea that has to be true to make an argument work. For example, when solving the pedigree part A problem, Martina stated a piece of evidence*, “Then the black circle would mean that they have it.”* Then she acknowledged an assumption *“and I also am assuming the circle is female.”* Martina solves the rest of the problem correctly, backing her claims with evidence.

Students acknowledged inferences when they recognized an inference in their reasoning. In the context of this study, an *inference* is a logical leap or bridge to a claim made with sufficient backing. For example, when solving the protein X problem, Sara made an inference, *“So I’m assuming those are more Van der Waals interactions”* which is backed by a piece of evidence, *“because there’s not, there’s not an oxygen or nitrogen or another electronegative atom sitting there.”* After making this inference Sara solves the problem correctly, backing her claims with rules.

Interestingly, though the function of Martina and Sara’s reasoning in these examples serve different purposes they both describe that they are *“assuming.”* Throughout the data set, students who produced solutions with HQR used the term “assume” or “guess” interchangeably to describe making an assumption or an inference in their thought process. Thus, instead of relying on key words to identify the use of “acknowledging assumptions” or “acknowledging inferences”, we had to distinguish inference versus assumption based on the functional role of the students’ metacognitive statement.

#### Identifying and orienting self to problem features

Students use self-questions to help them identify where something is in the diagram or to orient themselves to what they are looking at, which is associated with HQR. For example, after reading the protein X problem out loud Jeffery says, *“Okay, where is protein X?”* Then he answers his own question, *“Okay. It’s just the whole thing is folded into its tertiary structure surrounded by water molecules.”* Jeffery did not use any other metacognitive utterances in the rest of his thought process, but solved the problem correctly and backed his claims with rules.

#### Reassess the task before stating a claim

Students who use HQR restate the question in the problem in their own words before making a claim. For example, while nearing the end of solving the protein X problem, Eva asks, *“Um, do we predict that [the mutation] would affect the non-covalent interaction?”* Then, she responds with a claim, *“Yes, it would affect the non-covalent interaction, pointed to by the arrow.”* Eva solves the problem with mostly correct ideas (with a minor incorrect idea that didn’t impact the outcome of the problem), backing her claim with a rule.

### Purpose of Evaluating Utterances

#### Evaluating with self-questions to double-check accuracy, understanding, and what the question is asking

Evaluating utterances associated with HQR included three types of self-questions: self-questioning to evaluate accuracy, understanding, and what the question is asking. We found that there were very few examples of evaluative self-questions in our data compared to monitoring self-questions. Evaluative self-questions are different from monitoring self-questions because they occur when students assess their work after they provide an answer or conclusion to complete a phase of the problem-solving process.

Self-questioning to assess accuracy occurs when students ask themselves a question to appraise the correctness of a statement after describing their conclusion or answer. For example, while Renee was solving the pedigree problem part A, she stated the following claim to answer part of the question, *“So [individual] number two, right off the bat would have two ‘little a’s.’*” She then described a piece of evidence, *“but uh, we see that [individuals] three and four don’t have the trait”* to back the following claim, *“meaning that they either are heterozygous or they are homozygous ‘big A.’”* Then, she described a piece of evidence, *“Wait, both of which both genotypes would result in them not having the trait.”* Then she asks herself a **question to double-check the accuracy** of her previous ideas, *“But then if I were to give [individual] one a heterozygous genotype, that would mean that 50% of the offspring would have, right?”* and confirms, *“Yeah.”* Then, she described a piece of evidence, *“50% of the offspring would have the trait, and then 50% would be heterozygous.”* Renee successfully assessed the task and states, “*And whereas that is possible here because… Oh, it says the possible genotypes of the individual’s numbers one and two, I don’t need to necessarily choose between one genotype and the other.”* Renee solved the rest of the pedigree problem part A successfully, backing her claims with rules.

Only one student, Jeffery, asked himself questions to evaluate his understanding. Self-questioning to evaluate understanding occurs after “monitoring understanding” because it involves the assessment or appraisal of understanding. As mentioned previously (*Verification of knowledge accuracy and relevance: Monitoring with self-questions*), Jeffery solved the pedigree problem part B. Jeffery gave a solution with only correct ideas about almost all the individuals’ genotypes but monitored that he does not know what the second allele of individual two’s genotype is. The following paragraph elucidates how Jeffery identified the gap in his understanding using evaluative self-questioning.

After asking a self-question to check accuracy, Jeffery stated multiple correct claims about individuals one, three, and four backed by multiple pieces of evidence and a rule. Then, Jeffery monitored his understanding, *“And then individual number two, we know that they have to have at least one dominant…”* which is immediately followed by monitoring confusion, *“Okay wait, I’m getting confused. Yeah, so it’s X linked, so yeah.”* Jeffery stated a claim, *“So [individual] number two would have to have at least one dominant [allele].”* Then he monitors understanding, *“Not sure about the second [allele].”* This utterance is followed by a **self-question to evaluate understanding**, *“Are we sure about the second [allele]?”* followed by, *“Let me double check that because yeah, we’re not sure.”* Then, he described two claims*, “And then [individual] three would be ‘little a’ ‘little a’. And then [individual] four’s X would be ‘little a’”* He then monitored his understanding of his previous claim by backing with a piece of evidence, *“And we know that for sure because it’s not a shower or does not display [the trait].”* Then, he submitted the solution, indicating in his answer that the first allele was “big A” and he didn’t know what the second allele was. Even though he was not able to answer this part of the question, he was nearly there. Jeffery used both monitoring and evaluating skills to pinpoint what he understood and where there were gaps in his understanding; in this case the second allele for individual two’s second X chromosome. These data suggest that the ability to identify where and what the gaps are in your own knowledge is as or even more crucial for learning than the ability to identify what you already understand.

One student, Simone, used a self-question to evaluate what the problem is asking after describing a solution. As mentioned previously (*Retrieval of knowledge: Monitoring with self-questions*), while solving the protein X problem, Simone asked herself a conceptual understanding question to retrieve rules to back her claim. The following quote highlights how after retrieving the rules, Simone asked herself a question to evaluate what the question is asking. Directly after describing a rule that asparagine is a hydrophilic molecule, Simone asks, *“Well now that it’s hydrophilic it would also be folded different, but is that what the question is asking?”* Then, she rereads the problem out loud to evaluate the task and makes a correct claim to directly address what the question is asking, *“Yes, there would be a change in the type of non-covalent interaction.”* Simone solved the protein X problem successfully, backing her claims with rules.

### Illustrating Low-Quality Reasoning Responses during Problem Solving

Though this paper focuses on the types of metacognitive utterances associated with HQR, it is useful to compare this to the metacognition students used with low-quality reasoning (LQR). While problem solving, students would often use LQR because they did not know or remember the domain specific knowledge required to answer the questions from their previous learning experiences. For example, a student might not recall what non-covalent interactions are, which is an essential piece of knowledge for solving the protein X problem. If a student makes either a correct or incorrect claim without sufficient and relevant backing, it is considered LQR. The following examples from Quinn and Tabitha capture how students used LQR to answer the challenging biology problems.

While solving the pathway flux problem, Quinn first monitored familiarity, “*I’m trying to remember when I learned all this stuff. It’s been a while.”* Following this monitoring utterance, she remained silent for about 30 seconds until the interviewer prompted, *“What are you thinking now?”* Immediately after, Quinn stated the following correct claim as her final answer, *“I’m thinking that my answer is yes, it would affect it.”* Though her claim is correct, she did not provide any supportive backing, so this solution is considered LQR. Interestingly, after monitoring a lack of familiarity, Quinn does not think through the topics out loud. It is not clear whether she acts on her metacognitive monitoring. In contrast, other students such as Elaine monitored a lack of familiarity and then asked a self-question to recall knowledge, when solving the protein X problem (Table 3). When she wasn’t able to answer the question, she used a strategy to reflect on what she remembers which then sparked her memory and allowed her to recall the necessary knowledge to solve protein X problem with HQR (Table 3).

Tabitha, while answering the protein X problem, changed her answer multiple times and struggled to use sufficient backing outside of her own intuition to support her claims. While approaching her final answer, Tabitha restates the question in her own words, *“Do I predict that such a mutation would affect the non-covalent interaction pointed to by the arrow?”* Here Tabitha uses metacognitive monitoring in the form of self-questioning. Tabitha then answers her self-question, *“My gut instinct says no [the noncovalent interaction will not change] because of the nitrogen having three (electrons), but for some reason it makes me wanna say yes because then I think it could do four (bonds), but no again, because the carbon doesn’t wanna have four bonds. So regardless of what the nitrogen’s able to do, the carbon doesn’t want to bond with it there. So my instinct is to say no.”* Here, Tabitha expresses a lack of certainty about nitrogen and changes her mind multiple times; however, she is focused on the wrong idea needed to solve the problem. This example exemplifies how students may use metacognitive skills, such as self-questioning, to support problem solving but it does not always lead to high-quality reasoning if it is not productive, or does not advance problem solving.

### Metacognitive Utterances Associated with Low-Quality Reasoning

#### Monitoring

We found two monitoring utterances that were associated with students’ LQR responses. Students who used LQR to answer the biology problems often monitored a *lack* of familiarity and asked self-questions *without answering them out loud*. As a result, both of these monitoring utterances were unproductive to the problem-solving process. If a student monitors a lack of familiarity or recognizes that they do not remember the domain specific knowledge necessary to solve the problem, it is reasonable to expect their effort and metacognitive skills may only lead to a LQR response.

We found a plethora of self-questions associated with LQR and nearly all of them were not answered out loud by the student. For example, when solving the protein X problem, Michael states a piece of data*, “Okay, so leucine gets traded with asparagine.”* Then, Michael asks himself a self-question, “*And would [the mutation] affect the non-covalent interaction between those amino acids?”* He does not answer this question out loud and then follows up with another self-question where he attempts to monitor relevance, *“So my question is, um, is there even a, a reason to consider [amino acids] A, B, D and E, are those relevant?”* He does not answer either of these self-questions out loud throughout the rest of his response. Although Michael was able to monitor his gap in knowledge, he seems to lack the domain specific knowledge required to answer his own questions, and provided an incorrect and incomplete LQR response.

#### Evaluating

Some students who used LQR to answer the biology problems in our study evaluated their problem-solving strategies or the accuracy of their solutions. Evaluations made by students during LQR were often not productive for their problem-solving process. As an example, Erwin gave a LQR response for the pedigree problem part A; he unsuccessfully assessed the task and gave individuals 1-4 incorrect genotypes (though the problem asks only for individuals 1 and 2). Then he moved onto part B and read the question out loud to himself. After he finished reading, Erwin monitored his understanding with a self-question, *“What does it mean by X-linked?”* Then, Erwin ponders for a moment and describes, *“Hmm. X-linked… X-linked…. I feel like I kind of answered this because I kind of gave all [individuals 1, 2, 3, 4] genotypes [in part A].”* Then, he repeats his self-question again, *“So I guess for part B, X-linked dominant trait, what does that mean?”* He does not answer his question to remember what X-linked dominant means and follows up with how he will move forward *“I guess I would just label what I had in part A.”* Erwin then evaluated his part A solution, “*I didn’t realize that I [laugh] I didn’t have to give all of [the individuals genotypes] in my explanation.”* Following this evaluation, he typed his answer and describes how he adapted his answer from part A to answer part B and assessed that his answer was sufficient to submit, *“I added to my (part A) answer, so it could be a little different (for part B), but yeah, I’m okay with this.”*

When Erwin was typing his solution, he fell silent, and the interviewer prompted him to think out loud. Erwin then described his thought process for why he wanted to make the pedigree part B solution different from part A, revealing his misconceptions. He stated, *“My thought process was like trying to do all of the possible genotypes, which would include the shapes. And so, I worked out how I can add those in because it would be four letters instead of just two. And I needed to show that in my scientific explanation.”* Finally, as he submits the solution he evaluates the accuracy of his answer, *“So it might come off as a little weird, but I am pretty sure the outcome works out.”* Erwin’s data exemplifies how students can use frequent metacognitive utterances when problem solving, but their skills are unproductive if they lack accurate domain specific knowledge to answer their own self-questions to recall knowledge. Erwin was not able to evaluate effectively and successfully solve the problem because he was building his answer on a foundation of inaccurate scientific ideas.

#### Self-coaching found in LQR responses

We found multiple instances of self-coaching during LQR responses. Students use self-coaching to encourage themselves to keep solving after they face discomfort from monitoring a lack of understanding or prior knowledge (Halmo et al., 2024). In the absence of self-coaching, some students monitor lack of understanding or prior knowledge and then give up on problem solving (Halmo et al., 2024). For example, after reading the pedigree problem part A to herself, Chandra self-coaches, *“Okay, I’m gonna think out loud. I don’t have an answer but I wanna think out loud.”* Then, Chandra focused in on X-linked recessive traits, even though the question asks for the possible genotypes for individuals one and two if the trait is autosomal recessive.

Chandra described, *“Um, so if it’s a recessive trait and the mom has it, then her daughter could be a carrier. Like that’s the possible genotype. She could be a carrier but she won’t express it. I think just because like the dad doesn’t have it and she has two X chromosomes, but the son, he has like, I wish I could draw this out.”* The interviewer encouraged Chandra to draw out her thinking, and she attempted to draw a Punnett square for about a minute. Then, Chandra evaluated that her strategy was not effective, “*I’m not drawing [the Punnett square] ‘cause it’s not working.”* Chandra followed this evaluation with an incorrect guess, eventually leading to a LQR solution. In this example, Chandra used self-coaching to move past her discomfort of not knowing how to solve the problem. She continued problem solving by attempting the strategy of drawing a Punnett square. When this strategy didn’t work, she eventually stopped problem solving, yet she still submitted a solution. Though Chandra used self-coaching to initially move past discomfort, because she didn’t have the prior knowledge required to solve the problem, she provided a LQR solution.

#### Lack of prior knowledge is the main roadblock in LQR responses

As described previously, monitoring can aid students in the retrieval of knowledge, verification of knowledge accuracy and relevance, building of reasoning bridges, orienting themselves to what they are looking at, and reassessing what the question is asking. Evaluating aids students in double checking accuracy, understanding, and determining what the question is asking. Ultimately, metacognitive skills allow students to bridge the gap between their prior knowledge and specific features of a challenging problem, but if they lack sufficient domain specific knowledge required to answer, it is reasonable to expect that they may struggle to provide a solution backed with HQR.

### Individual Differences in Metacognition Use

Some students displayed more sophisticated and unique metacognitive skills that led to higher quality reasoning in comparison to their peers in the study. Simone, Jeffery, and Renee all used evaluative self-questions and multiple monitoring utterances in their think aloud solutions. These three students consistently displayed the highest quality reasoning across the three problems. In addition, Kathleen, Elaine, and Sara, though they didn’t use any evaluative self-questioning, displayed a range of monitoring skills associated with high-quality reasoning. It is possible that these students were not challenged enough to use evaluation or it is not a skill that they have developed to use during problem solving. These six students, when compared to the other 33 students, stand out as displaying a more sophisticated ability to generate high-quality scientific reasoning with metacognition to solve the problems.

To illuminate some more of the unique individual differences, we highlight some think aloud data from Renee. To help her get further in the pedigree part B problem. Renee used a strategy of making an educated guess. Many students in our study made guesses when they were stuck or they couldn’t recall a piece of knowledge that was required for solving the problem. The guesses made by students were almost always incorrect and were not productive for the problem-solving process. Interestingly, only one student, Renee, was able to successfully use a partially correct guess as a strategy that led her to a completely correct solution, where she backed her claims with rules.

After reading the pedigree part B question, Renee first evaluated difficulty, *“Okay, so [sighs] if this is X-linked dominance, gosh, X-linked dominant, if it were Y-linked, it would be an easier question*” Then she described a piece of evidence, “*X-linked dominant trait, meaning that the mom and the dad have it.”* Renee monitored familiarity and understanding of a piece of evidence, “*but I’m trying to remember what that really means because I know that a Y-linked genotype would just mean that the dad is always gonna have it, but here, the, the mom has it.”* Next Renee improvised by making a guess, “*So, I’m just going to use the power of inference [laughs] and say that it kind of works like autosomal traits.”* Though her guess is not definitively correct, for the context of answering the problem it was correct enough to move past the gap in her understanding. Ultimately, by making this guess, Renee was able to provide complete and correct answers for all four individuals’ genotypes, backing her claims with rules.

Our data suggest that some metacognitive skills, such as monitoring with self-correction, are more automated for students. To highlight the automaticity of self-correction, we describe an example from Renee’s pedigree part B solution. In Renee’s think aloud interview, she corrected her own internalized thoughts before they are fully externalized into words. The speed of her internal monitoring seems to be faster than the speed of her speech. While solving the problem, she first asked herself a question to recall knowledge, *“What was I gonna say about individual two?”* Then, Renee stated a claim, *“I’m gonna say X ‘big A’ and then X ‘little a’, and then X…”* She immediately self-corrected and states another claim, *“No, there’s no way it can be ‘big A’ ‘big A.’”* Interestingly, in her first claim she never said that individual two’s genotype could X ‘big A’ ‘big A’ but we reason that she was beginning to say that idea (*“and then X…”*) before she self-corrected. Then, Renee described a piece of evidence to support her self-corrected claim, “*because if it were ‘big A big A’, it would mean that the son would inherit the Y from his dad, but then an X with a ‘big A’ from his mom no matter what. So, he would have the trait.”* Finally, she stated a correct claim about individual two’s genotype, “*I’m just gonna say that the only possibility is that it’s ‘big A’ ‘little a.’”* As mentioned previously, Renee was able to solve the problem successfully, backing her claims with rules.

### Self-questions Are the Engine of Metacognition During Problem Solving

Self-questioning during individual problem solving is very important for generating high-quality reasoning but it is only productive when a student accurately answers their own question out loud. Across the students who used metacognition to solve the problems with HQR, we found that self-questions often were the result of or produced other metacognitive utterances. Self-questions were also useful at any point in the problem-solving process: from initially reading the problem to submitting a final solution. Throughout our data, self-questions seemed to act as a propeller to guide students to further monitor and evaluate their thinking. If answered correctly out loud, self-questions can be used to monitor and evaluate students’ prior knowledge and build sophisticated scientific arguments to answer biology problems.

In some way self-questions are analogous to a car engine and prior knowledge represents the fuel. When a car runs out of fuel, it stops because it can no longer drive any further. Similarly, when a student runs out of the necessary prior knowledge, they cannot solve the problem. They may attempt to rev the engine by using a self-question, but may not be successful, because they have run of prior knowledge. On the other hand, a student who has sufficient prior knowledge, or enough fuel in their tank, can use self-questions to rev their engine and propel them to their final solution. We found a variety of monitoring and some evaluative self-questions displayed by the students to generate high-quality reasoning. Asking and answering self-questions is a critical part of life science students’ problem solving that can lead to more metacognition, propelling the students further in their problem-solving process.

## DISCUSSION

When we asked fourth-year life science students to solve challenging biology problems out loud, they used a variety of monitoring, some evaluating, and nearly no planning utterances associated with HQR. We were able to identify how metacognitive regulation (often self-questions) during problem solving provides links to bridge the gap between a student’s prior knowledge and specific features of a challenging problem. Here, we aim to expand metacognitive theory by providing empirical support for the use of automated metacognition during problem solving. Rooted in our student think-aloud data, we propose and characterize and a model of metacognitive phases during problem solving. Finally, we share a metacognitive self-questioning scaffold that can be used to guide and facilitate metacognition during individual and group problem solving. We reason that metacognitive interventions to promote the use and integration of planning, monitoring, and evaluating skills are necessary. We can help life science students’ access, utilize, and leverage their prior knowledge from previous learning experiences through metacognitive interventions, such as a self-questioning scaffold, to enhance their HQR skills for productive problem solving.

### Monitoring dominates over evaluating and planning skills during HQR

We found that fourth-year life science students used a variety of monitoring skills but do not readily use planning and evaluating skills during problem-solving that leads to HQR. Evaluating utterances were less common than monitoring utterances. Thus, without intervention, many life science students have developed some monitoring skills but did not show much evidence of planning or evaluating during think aloud problem solving. This aligns with our prior work where we used a think-aloud protocol to investigate the types of metacognitive regulation that first-year life science students used while solving biology problems (Halmo et al., 2024). First-year students used a variety of monitoring, engaged in limited evaluating, and lacked planning skills. We also found that first-year students use self-coaching to encourage themselves to keep solving after they face discomfort from monitoring a lack of understanding or prior (Halmo et al., 2024). Without self-coaching, some first-year students monitor lack of understanding or prior knowledge and then give up on problem solving (Halmo et al., 2024).

Interestingly, we did not find any evidence of fourth-year students using self-coaching when solving challenging life science problems with HQR, but some participants used self-coaching when giving LQR responses. Lack of prior knowledge was the main roadblock for students who gave LQR responses compared to HQR responses. If a student lacks the prior knowledge required to provide an answer with HQR, it is reasonable to expect that they will face some discomfort during problem solving. During the discomfort they can either use self-coaching to push themselves further in the problem-solving process or give up. In our data, we saw that some students used self-coaching in LQR responses to push themselves further, however, in the end they often stopped problem solving, because prior knowledge was the main determinant of their problem-solving outcome.

Results from Versteeg et al. (2021) also align with our previous and current findings of metacognitive regulation skills used to solve scientific problems. Versteeg et al. (2021) utilized a think aloud protocol to capture in-the-moment metacognition skills when undergraduate, preclinical medical school students in the Netherlands solved conceptual physiology problems. Echoing the outcomes of this current study, they also found that less time was devoted to unprompted planning and evaluating as compared to various forms of monitoring (Versteeg et al., 2021). The researchers also described that even when they prompted students to evaluate their thinking, they spent little time evaluating (Versteeg et al., 2021). However, unlike the current study, they did not distinguish the types of metacognitive regulation skills that actively advanced students’ problem-solving process and led to solutions backed with HQR.

### Why are only some students using evaluating skills during problem solving?

Other studies have indicated that students evaluate their peers’ ideas more readily than their own ideas (Halmo et al., 2022; Bremers et al. 2025). As such, students can benefit from peers evaluating their thoughts during group work; as they may not spontaneously engage in evaluation while working individually (Chiu & Khoo, 2003; Rickey & Stacy, 2000). In the context of group work in an upper-division biology course, we previously conducted a study to investigate the types of social metacognition that students engage in when constructing HQR during problem solving (Halmo et al., 2022). Social metacognition can be defined as awareness and regulation of others thinking for the purpose of learning. In units of HQR, we found that students often posed questions to their peers, such as questions to monitor understanding or request information; which allowed other students to answer the question and evaluate or correct their ideas. We also found that evaluative questioning was a key feature in constructing HQR units during group work, but only if another peer answered the question.

The results from our study also indicate that individuals who use HQR during problem solving ask themselves questions and answer them out loud to monitor and evaluate their ideas during problem solving. Unlike social metacognition, if an individual asks themselves a question and does not know the answer, they may not be able to solve the question without consulting an outside resource. When working in groups, students’ can engage in social metacognition by posing questions that they can’t answer themselves to their peers and receive an answer. If group problem solving is occurring in a classroom setting, students can also ask questions and practice social metacognition with their instructor or teaching assistant. We reason that in both individual and social metacognitions contexts, posing questions to yourself or others, and receiving an answer is a critical aspect of using metacognition to navigate the construction of HQR during problem solving.

We also found that a small subset of students engaged in more sophisticated metacognitive skills that led to higher quality reasoning as compared to their peers, thus illuminating individual differences across our data set. Three out of 39 students who engaged in multiple types of productive monitoring and evaluating across all three problems consistently displayed the highest quality reasoning. The productive evaluating utterances that lead HQR were evaluative self-questions, which were scarcely used by other students. Students who used sophisticated metacognitive utterances to solve problems did not show evidence of planning, like the rest of participants in the study.

### Why aren’t students using planning skills during problem solving?

During problem solving, planning is used to select strategies, determine how and when to use them, and make predictions about the solution (Stanton et al., 2021). Across the data set, regardless of reasoning quality, we found a lack of planning used during problem solving. It is possible that this was the result of the problems we used being well-defined (also referred to as well structured), which means they state clear givens, goals, and have one solution (Ge & Land, 2003; Jonassen, 2000). The problems students solved were also straightforward, and short enough that they did not require students to plan in order to solve the problem effectively (Halmo et al., 2024).

It is important to consider that real-world problems are more likely to be ill-defined than well-defined. Problems are ill-defined (also referred to as ill structured) when they lack a clear end goal and can be solved with more than one accurate solution (Ge & Land, 2003; Jonassen, 2000). Ill-defined problems require more extensive planning skills for solving as compared to well-defined problems. As researchers and practitioners, we aim for students to develop field specific wisdom and metacognitive skills that they can transfer into graduate school or future careers; where they will certainly take on real world, ill-defined problems. We speculate that if we give students a more ill-defined task, it may elicit planning skills.

The amount of domain-specific knowledge that a problem-solver has impacts the types and amount of time used for planning (Davidson & Sternberg, 1998). Effective problem-solvers with domain-specific expertise often spend time classifying the question based on their deep conceptual understanding, and they spend more time planning than novices (Chi et al., 1981; Davidson & Sternberg, 1998). Instead, novices use little or no planning, classify questions with surface level features, and jump straight into solving for the solution without making predictions or determining the path they will take and strategies they will use. (Chi et al., 1981; Davidson & Sternberg, 1998). As a result, novices are more prone to making mistakes and providing incorrect solutions (Chi et al., 1981). More research is needed to understand how planning skills are impacted based on how well-defined the problem is and the domain specific knowledge of the problem solver.

### Support for automated metacognition during problem solving

Metacognition researchers have continued to contemplate the existence of automated metacognitive processes and how to measure them (Azevedo, 2020). Some researchers argue that automated metacognition is an unconscious process while others reason that metacognition is conscious by nature, so automated metacognition refers to conscious thought that occurs with little attention or effort (Azevedo, 2020; Reder, 1996; Samuels et al., 2006). However, both sides of the argument have merit, as automated metacognition may be more nuanced. It is likely that metacognition encompasses both unconscious and conscious processes (Timmermans et al., 2012). With our think-aloud data collection method, we assume that we are only able to capture the conscious automated metacognition, requiring little attention or effort, since we are directly accessing a student’s conscious working memory (Ericsson & Simon, 1980).

Veenman (2015), describes how conscious, effortful metacognitive processes can become automated over time with sufficient practice. Unlike planning and evaluating, monitoring skills may be consistently running in the background, detecting errors or anomalies as they occur in real time (Veenman, 2011b, 2015). Though monitoring skills can be automated, most metacognitive skills are not practiced enough and may never become fine-tuned, fast, and effortless (Veenman, 2011b, 2015). Thus, in this study, monitoring skills such as self-correction, or when a student corrects themselves while talking out loud from incorrect to a correct idea, may be more automated as compared to other skills. We found that when students monitored an error in their speech and self-corrected to fix the mistake, it was often fine-tuned, fast, and effortless. We also hypothesize that monitoring with self-correction can be more widely used across tasks compared to other domain-specific metacognitive skills.

One student from our study, Renee, exemplifies what automaticity of monitoring with self-correction might look like while solving the pedigree problem part B. When she was solving for individual two’s genotype, she stated one claim and began to state another, *“I’m gonna say X ‘big A’ and then X ‘little a’, and then X…”* She immediately self-corrected and made another claim, *“No, there’s no way it can be ‘big A’ ‘big A.’”* We noticed that she did not say individual two’s genotype could be X ‘big A’ ‘big A’ out loud, but we reason that she was beginning to say that idea (*“and then X…”*) before she corrected herself. Then, Renee described a piece of evidence that provides a new correct claim for individual two’s genotype, and solved the rest of the problem with correct solutions, backed with HQR. In this example, it seems that Renee self-corrects her own internalized thoughts before they are fully externalized into words. The speed of her internal monitoring is faster than the speed of her speech, which we reason could be evidence of monitoring automation. Our study goes beyond mere contemplation of automated metacognition existence and as far as we are aware, provides the first evidence that monitoring with self-correction may be more automated than other metacognitive regulation skills.

### The impact of students’ self-questions during learning tasks

When students ask questions, it demonstrates that they are engaged in the learning task and are attempting to integrate the new knowledge presented to them with what they already know (Chin & Osborne, 2008). Problem solving requires students to integrate any new information from the problem with their own prior knowledge (Frey et al., 2022) and question posing may help guide their thinking. As displayed by the students in our study, asking and answering self-questions out loud is a critical part of life science students problem solving. Throughout our data, self-questions seemed to act as an engine to guide students to further monitor and evaluate their thinking and helped them build sophisticated scientific arguments to answer biology problems.

To the best of our knowledge, the finding of life science students using self-questions during problem solving is unique, and no prior study has demonstrated this specific outcome before. Yet, the concept of using self-questions as a strategy for learning tasks is far from being a novel one. Self-questioning during a learning task, like reading, is a highly metacognitive skill, as it requires a student to monitor and identify what they do and do not know (King, 1989). As such, there is a large body of literature spanning multiple decades that indicates that self-questioning is an effective strategy to improve reading comprehension for kindergarteners to college students and for students with and without learning disabilities (Joseph et al., 2016; Wong, 1985).

In addition to aiding in reading comprehension, self-questioning has been investigated as a strategy to improve comprehension during and after lectures for high school and college students. For instance, King (1989) describes that by asking self-questions during a lecture, students enhanced their understanding during class. Similarly, another study strategically trained high school students how to self-question with prompts during lecture significantly increased student comprehension (King, 1991a). Even when the prompts were removed, students maintained the self-questioning strategy and retained the information they learned from lecture for longer than students who did not use the strategy (King, 1991a). Self-questions have also been found to be a successful post-lecture study strategy to help college students retain information for longer than reviewing or summarizing notes (King, 1992).

Although not studied in college settings, a few studies have explored how guided questions can help elementary students’ problem solve more effectively. For instance, third-grade students who encountered difficulty with subtraction and addition word problems were able to more effectively read and solve the problems when trained how to use a self-questioning scaffold (Sulaimon et al., 2025). The positive effects on their reading comprehension and problem-solving abilities were consistent even when the scaffolding was removed (Sulaimon et al., 2025). Another study investigated how the use of a problem-solving prompt card (also called a regulatory checklist), which included a series of planning, monitoring, and evaluating questions, impacted fifth grade students’ problem-solving abilities when working in pairs (King,1991b). The authors found that pairs of students who used the prompt card while problem solving had enhanced performance, but only when they were adequately guided and trained how to use the strategy (King, 1991b).

Based on the findings from King (1991b), Schraw (1998) recommended the use of a regulatory checklist of questions as an effective problem-solving strategy for promoting metacognitive awareness. It is important to note that the questioning strategy in King (1991b) is peer-questioning, and not self-questioning. However, Pate & Miller (2011) adapted the peer-questioning regulatory checklist from King (1991b) into a self-questioning regulatory checklist that high school students were instructed to use while solving electrical circuit theory questions. They found that students who received explicit training and modeling on how to use the self-question regulatory checklist during problem solving performed significantly higher than students who did not use the checklist (Pate & Miller, 2011).

Prior research on using self-questions as a learning strategy is largely focused on reading and lecture comprehension, with few studies examining its use for problem solving activities. Based on our findings, asking and answering self-questions is a critical metacognitive skill that life science students use spontaneously in problem solving with HQR. Thus, we reason that modeling, training, and prompting life science students to ask and answer self-questions may be an effective intervention to enhance metacognitive regulation skills during problem solving. Educators must help students understand that question-posing lies at the heart of scientific inquiry; driving the problem-solving process that advances scientific progress.

### Metacognitive phases during problem solving with HQR

Based on our data, we propose three different phases of metacognitive regulation for problem solving with HQR: monitoring, reasoning, and evaluating. We also hypothesize a planning phase, which was not found in our student data but if it occurs, it will appear before the previous phases. Thus, making the HQR problem-solving phases: **planning, monitoring, reasoning,** and **evaluating** which are facilitated through self-questioning (Fig. 1). We want to emphasize that we intentionally chose the word “phase” over “stage” because the problem-solving process to build a solution with HQR is highly iterative and recursive, not linear or sequential. Additionally, since certain monitoring processes are often more automated than evaluating and planning skills (Veenman, 2011, 2015) they can run in the background to detect errors and anomalies. Thus, across all problem-solving phases, students can monitor accuracy of their knowledge and use self-corrections to verify and correct in real time if they monitor an error in their understanding or thought process.

**Fig. 1.**
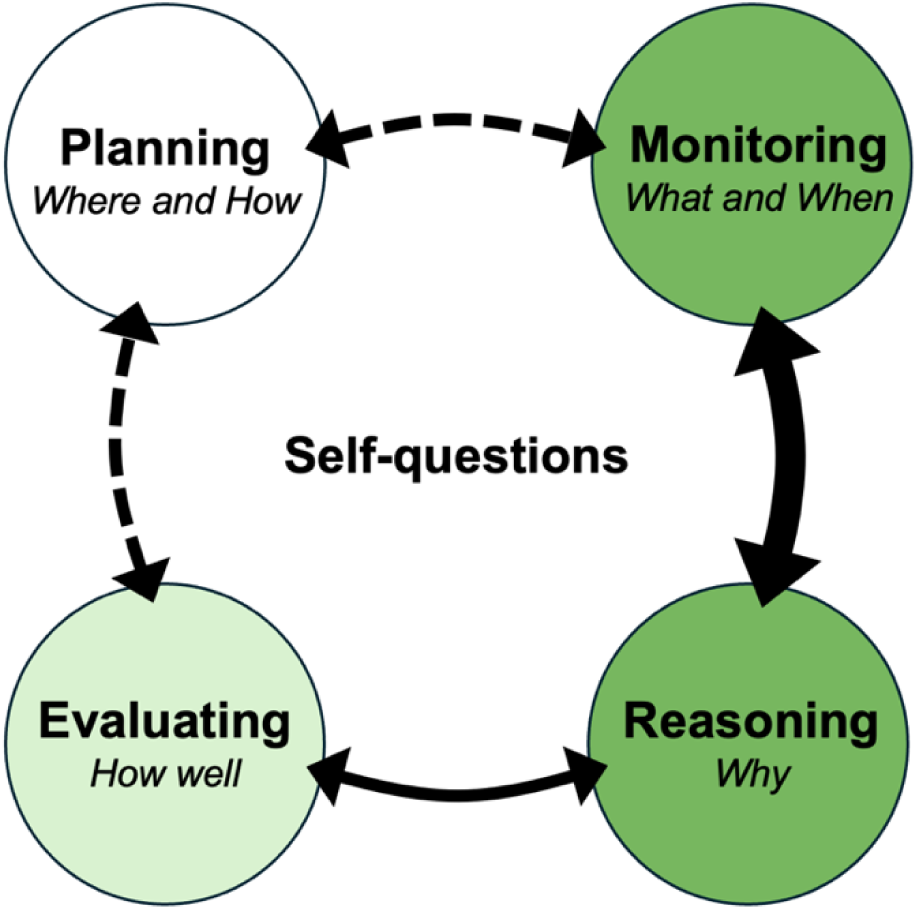
Proposed metacognitive problem-solving phases during high-quality reasoning (HQR). Self-questions can help generate and guide student thinking through the planning, monitoring, reasoning, and evaluating phases. We hypothesize that if given a sufficiently challenging, ill-structured question, an expert problem solver will begin solving at the planning phase, followed by monitoring, reasoning, and finally the evaluating phase. The dotted arrows indicate hypothesized relationships and solid arrows denote empirically supported relationships between phases. We propose that if the solver reaches the evaluating phase and evaluates that their solution is incorrect, insufficient, and/or doesn’t make sense, they may have to revisit the planning phase. Then they can adjust their plans and restart the cycle with updated information so they can provide a successful solution. If the solver runs into a roadblock, such as an error in their solution, they may have to go backwards in the loop, to correct their mistakes and continue the problem-solving process. Students in our study often utilized the monitoring and reasoning phases (indicated by dark green), a few ventured into the evaluating phase (light green), and none used the planning phase (uncolored). The monitoring and reasoning phases are closely tied together, as participants who gave HQR solutions often moved back and forth between them multiple times.

While here we are studying metacognition, which is the awareness and control of one’s thinking for the purpose of learning, we also want to recognize the role of cognition in metacognition. Cognition is the mental action of processing information. Thus, the phases below, though largely metacognitive, are also accompanied by cognitive processes. For example, when solving the protein X problem Jeffery asked himself a question (metacognitive) *“Okay, where is protein X?”* Then he answers the questions out loud (cognitive), *“Okay. It’s just the whole thing is folded into its tertiary structure surrounded by water molecules.”* Ultimately, in this study, we used think-aloud data to access the complex cognitive and metacognitive processes that students use while building the “physiology” of their scientific argument (Toulmin, 1958/2003). As such, the phases described below integrate both cognitive and metacognitive processes to build a model of metacognitive phases during problem solving with HQR.

#### Planning Phase (Where and How)

We hypothesize that a problem-solver plans for where they will go in their problem-solving process by making predictions about the solution or what steps to take. They also determine how they are going to solve the problem by making a plan and selecting strategies. This phase also includes assessing the task of the problem. As they are solving, they can monitor and adjust their plans and predictions as necessary.

#### Monitoring Phase (What and When)

Students access and search their knowledge for what information they need to solve the problem. As prior knowledge is the main determinant of a participant’s performance, accessing the knowledge available from memory is an essential step in the problem-solving process. When building HQR solutions, students may monitor their understanding and familiarity, and by using self-questions to aid in the retrieval of knowledge. After retrieving the information from memory, students must monitor the relevance of the knowledge; or when is appropriate to use this piece of knowledge, based on the conditions of the problem. This phase can include monitoring if there are aspects of the problem that are relevant or irrelevant. We reason that when solving real world problems, solvers can and should access external knowledge resources (peer reviewed literature, experts in the topic, etc.) from beyond their own prior knowledge in the monitoring phase.

#### Reasoning Phase (Why)

In the reasoning phase, a student provides a piece of data, evidence, and/or a rule, for why they can make an inference or claim. This is where a student builds their scientific argument with appropriate backing and creates reasoning bridges to solve the challenging problem. Though not always required during problem solving, students can also acknowledge their assumptions and inferences in the reasoning phase. We define an assumption as a provisional premise or idea that must be true to make an argument work. An inference is a logical leap or bridge from a premise to a claim made with sufficient backing.

#### Evaluating Phase (How well)

Though rare in our data, students evaluate with self-questions to double check accuracy, understanding, and what the question is asking while forming a HQR response. They check for how well their scientific argument is made and double check if there are any mistakes or misunderstandings before they submit their solution. In this phase, students ensure that they understand their solution, and that their solution answers the problem and is correct.

In our study students used metacognition to retrieve *what* knowledge they need, *when* they need to use it, *why* they can use the knowledge to make inferences and build solutions, and *how well* their scientific argument is made before they submit. We largely found that students used a variety of monitoring skills and navigated the **monitoring** and **reasoning** phases, with only a few utilizing the **evaluation** phase, and limited evidence of the **planning** phase. Our data suggest that without intervention, some students may not fully develop their monitoring skills. We reason that metacognitive interventions to promote the use and integration of planning, monitoring, and evaluating skills are necessary to help aid students to further develop HQR skills.

### Implications for Instructors

Based on the results of this study, we provide multiple implications for instructors. First, we propose that students may benefit from thinking out loud during problem solving since there is an explicit memory benefit from producing speech (Forrin et al., 2012; Forrin & MacLeod, 2018). By verbalizing thoughts, students have to actively engage and may be compelled to monitor and evaluate their thinking instead of relying on intuition or feelings when coming to a solution. Thus instructors should encourage students to think out loud while problem solving by themselves outside of class. Students can also practice thinking out loud by collaborating on group problem solving activities during class time. By working with peers, students have the opportunity to engage in social metacognition, which includes monitoring and evaluating each other’s thoughts with questions during problem solving activities (Halmo et al., 2022) in addition to individual metacognition.

Here, we propose a self-question scaffold to promote domain specific metacognitive regulation skills and to facilitate the proposed problem-solving phases: planning, monitoring, reasoning, and evaluating (Table 5). In the context of undergraduate chemistry courses, implementing a metacognitive scaffold, such as Goldilocks Help workflow has been found to support students’ planning and evaluating skills for effective problem solving (Vo et al., 2024, 2025). Our proposed scaffold is rooted in student data, as we adapted the types of self-questions that our participants asked and answered out loud when building a solution backed with HQR. In our study, we found that students used a variety of monitoring skills and navigated the monitoring and reasoning phases. We only saw some students utilizing the evaluation phase and limited evidence of the planning phase in our data, pointing to the importance of explicitly guiding and prompting students to evaluate and plan. This intervention may help life students practice their metacognitive regulation and reasoning skills while solving challenging problems.

**Table 5.** Metacognitive self-question scaffold for problem solving.

| **Problem Solving Phase** | **Self-Questions** |
|---|---|
| **Planning** | What is the question asking me to do?<br>What predictions can I make about the solution (if any)? |
| **Monitoring** | What do I know/remember or not know/not remember about the information given in the problem?<br>What aspects of my knowledge or the prompt are relevant or irrelevant? |
| **Reasoning** | What assumptions, inferences, or claims can I make based on my own knowledge and the information given in the question?<br>Do my inferences and claims have adequate scientific backing? |
| **Evaluating** | Does my answer make sense and is accurate?<br>Did I answer what the question is asking me? |

The most effective way to employ metacognitive interventions for learning gains is to integrate the scaffolding directly into the structure of the problem-solving assignment (Kramarski & Friedman, 2014; Veenman, 2011b; Volet, 1991). Thus, we recommend that instructors introduce the self-questioning scaffold to their students by integrating into a problem-solving activity (see Supplementary Information for an example). Additionally, instructors should model how to think aloud, coach students on the step-by-step process, and provide explicit reasoning for where, when, and why they should utilize the self-question scaffold (Volet, 1991). When students become more comfortable using the self-questioning strategy, the scaffolding can be removed, also known as scaffold fading (Belland, 2011). After students receive adequate guidance and practice enough to feel comfortable using the strategy, we aim for the metacognitive self-question scaffold to be used as a reference to guide and facilitate metacognition during individual or group problem solving. Through the use of this self-question scaffold, we can prompt students to self-question and use other various metacognitive skills to solve the problem with strong scientific backing.

## CONCLUSION

Ultimately, this study demonstrates that students who ask and answer self-questions to monitor and evaluate their thinking build strong solutions backed with high-quality reasoning (HQR). We found that self-questioning is a critical skill that can lead to more metacognition, propelling students further in the problem-solving process. Self-questioning and other metacognitive regulation skills provide links to bridge the gap between a student’s prior knowledge and specific features of a challenging problem. After conceptualizing a model of metacognitive phases during problem solving with HQR, we created a self-question scaffold. Future research should investigate how effective the proposed self-question scaffold is for improving students’ metacognitive regulation and reasoning skills during problem solving. As educators, we can use interventions to equip students with metacognitive regulation tools for effective problem solving. To help our students to find the right answers, we have to teach them how to ask the right questions.

## Supporting information

Pedigree Problem and Self-Questioning Scaffold

## ACKNOWLEDGEMENTS

Thank you to the College Learning Study participants who made this research possible; we are very grateful for the opportunity to learn from them. We also thank the College Learning Study research team for their effort, dedication, and support on this project.

This work was supported by the National Science Foundation (NSF CAREER 1942318). Any opinions, findings, and conclusions or recommendations expressed in this material are those of the author(s) and do not necessarily reflect the views of the National Science Foundation.

