## Supplementary material for "From Problems to Solutions: How Self-Questions Drive Strong Scientific Reasoning": Pedigree Problem and Self-Questioning Scaffold

**Supplementary Information**

**Pedigree problem (Part A and B) used during data collection**

The pedigree below could represent an autosomal recessive trait or an X-linked dominant trait. Each individual is labeled so you can refer to them by their number. In your answers, use the letters A ("big A") and a ("little a").

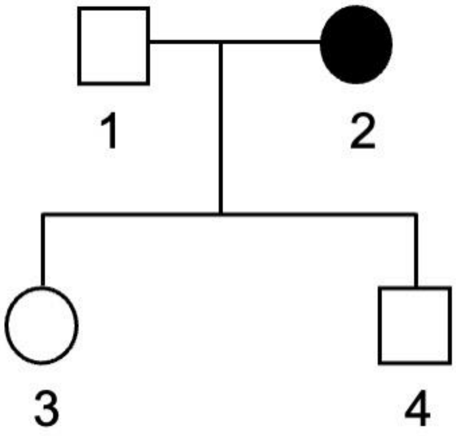

A) If this is an autosomal recessive trait, what are the possible genotypes of the individuals with numbers 1 and 2? Provide a scientific explanation to support your answer.

B) If this is an X-linked dominant trait, what are the possible genotypes of the individuals with numbers 1, 2, 3, and 4? Provide a scientific explanation to support your answer.

**Example of how to integrate the self-questioning scaffold into a problem-solving activity**

**Pedigree Problem**

The pedigree below represents X-linked dominant trait. Each individual is labeled so you can refer to them by their number. In your answers, use the letters A ("big A") and a ("little a").

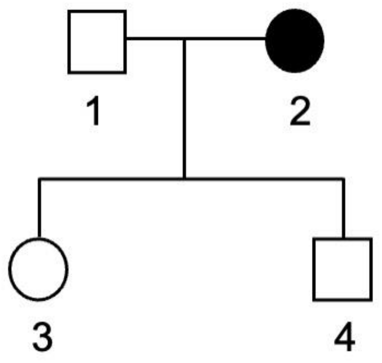

If this is an X-linked dominant trait, what are the possible genotypes of the individuals with numbers 1, 2, 3, and 4? Provide a scientific explanation to support your answer.

**Before**, **During**, and **After** solving, ask yourself the following questions out loud and briefly jot down your answers.

**Planning: Before solving**

- What is the question asking me to do?

- What predictions can I make about the solution (if any)?

**Monitoring: During solving**

- What do I know/remember or not know/not remember about the information given in the problem?

- What aspects of my knowledge or the prompt are relevant or irrelevant?

**Reasoning: During solving**

- What assumptions, inferences, or claims can I make based on my own knowledge and the information given in the question?

- Do my inferences and claims have adequate scientific backing?

| **Solution:** |
| --- |

More on the next page ☺

After writing your solution, ask yourself the following questions and briefly jot down your answers. You can adjust your previous solution in the updated solution box below or leave it blank if you do not have any changes from your solution box above.

**Evaluating: After solving**

- Does my answer make sense and is accurate?

- Did I answer what the question is asking me?

| **Updated Solution:** |
|---|
